# A Multi-omics Study on the Oncogenic Roles and Clinical Significance of Dynactin Family Gene (*DCTN1-6*) Expression in Liver Hepatocellular Carcinoma

**DOI:** 10.1101/2022.06.18.496693

**Authors:** Md. Asad Ullah, Tahani Tabassum, Afrah Rashid, Nafisa Nawal Islam, Moon Nyeo Park, Abu Tayab Moin, Bonglee Kim

## Abstract

In this study, we employed a comprehensive database mining approach to examine the possible oncogenic roles and clinical relevance of Dynactin family genes (*DCTN1*-6) in Liver Hepatocellular Carcinoma (LIHC). All the *DCTN*s were observed to be differentially expressed in LIHC tissues compared to the adjacent normal liver tissues. Most of the *DCTN*s were discovered to be aberrantly methylated (less methylated) and contain multiple somatic mutations (alteration frequency: 0.2-2.5%) in LIHC tissues. Overexpression of *DCTN*s was mostly associated with poor overall and relapse-free survival of LIHC patients. Alongside, all the *DCTN* genes were reported to be overexpressed across different demographic and clinical conditions, i.e., age, cancer stage, tumor grades, and metastatic stages of LIHC patients. *DCTN* expression was also associated with the infiltration levels of different immune cells, i.e., B cell, T cell, and macrophages in LIHC microenvironment. The co-expressed genes of *DCTN*s in the LIHC tissues were previously found to be involved in oncogenic processes in different cancer types and control crucial biological processes, i.e., nucleotide metabolism, RNA degradation, and chromosome organization. Later, the expression pattern of *DCTN*s was validated in two independent microarray datasets (i.e., GSE17856, GSE98383), which also supported our initial findings. All these findings suggest that *DCTN*s and their transcriptional and translational products are potential prognostic and therapeutic targets for LIHC diagnosis and treatment. This study will help further the development of *DCTN*-based diagnostic and therapeutic measures for LIHC and translate them into clinical implications.

## 1. Introduction

Primary liver cancer is the sixth most commonly diagnosed cancer and one of the leading causes of cancer-related malignancies worldwide. With an estimated 841,000 cases and 782,000 deaths in 2018, liver cancer ranked 7^th^ and 3^rd^ amongst different cancer types in terms of global incidence and mortality, respectively. Liver Hepatocellular carcinoma (LIHC), referred to as hepatoma, is the most prevalent histologic primary liver cancer encompassing around 75% of all liver cancer malignancies [1]. The burden of LIHC varies markedly due to a variety of factors, including demographic characteristics such as sex, age, ethnicity; hepatitis B or hepatitis C infection; liver flukes prevalent in endemic areas; behavioral factors such as consumption of tobacco and alcohol; metabolic syndromes such as obesity and diabetes; dietary factors; aflatoxin exposure; and genetic susceptibility [2–4]. This dominant histologic type of cancer mostly prevailed across Eastern and Southern Asia, middle and western Africa, Melanesia, and Micronesia [5]. However, in recent years, the rates of LIHC have drastically abated across some other regions of the world [6–8], raising major global health concerns. Since HBV and HCV are considered to be the major etiological causes of LIHC in Asia and Africa [9,10], implementing mass HBV vaccination in these countries rather than therapeutic advancements may account for the decrease in newly diagnosed cases of LIHC.

LIHC is typically diagnosed at the advanced stage when patients have already experienced some degree of liver impairment to become symptomatic, and unfortunately, there is no fully effective treatment as of today that has assured improved survival on administration against LIHC. Orthotopic liver transplantation (OLT) is considered the best therapeutic option for LIHC, but unfortunately, it is not suitable for advanced-stage LIHC or underlying liver cirrhosis [11]. Currently, other therapeutic approaches such as radiofrequency (RF), microwave ablation, subsume chemoembolism, radioembolism, and systemic treatments with sorafenib and regorafenib are in practice, but despite all these advancements, the complete cure of advanced-stage liver cancer remains a grey zone [12–14]. Moreover, the mortality rate of LIHC associated with therapy is, unfortunately, very high [15]. Consequently, genomic targets and assessment of the genetic basis of LIHC have garnered major attention in recent years to target major molecular pathways associated with LIHC for therapeutic intervention.

Dynactin (*DCTN*), a conserved, ubiquitously expressed, large multiple-subunit protein complex, is an essential cofactor for the microtubule motor cytoplasmic dynein-1 [16]. The protein comprises six distinct subunits referred to as dynactin 1-6 (*DCTN* 1-6), and all of these subunits are crucial for the structure, conformation, and function of the entire protein complex [17–19]. Unlike most large protein complexes, the dynactin molecule has a highly unusual subunit stoichiometry. The core protein structure is divided into three major structural domains, the sidearm shoulder, the actin-related protein Arp1 filament, and the pointed end complex [20], and the filament is nine subunits long consisting of two protofilaments [21]. Most of the *DCTN* structures are involved in the interaction of a variety of cellular structures, as well as the activation of most of the eukaryotic cytoplasmic proteins dynein, dynein cargo adaptors, kinesin-2 motors, and microtubules [20,22]. The dynactin-dynein complex is involved in crucial cellular and axonal transport, as well as, regulates the efficiency of fast retrograde transport of vesicles and organelles in cells [21]. Besides, dynactin plays a vital role across the cell division, such as mitotic alignment, microtubule-kinetochore attachment, nuclear envelope breakdown, maintaining centromere integrity, and spindle organization [23–26]. The undeniably significant role of dynactin in maintaining a healthy cell cycle and interactions with a wide range of cellular structures, particularly those involved in dynein movement, raises major concerns regarding the probable association of this gene expression profile with distinct cancer types.

Several studies have recently demonstrated the association of *DCTN* genes and different *DCTN* protein subunits with several cancer types. Studies reported that *DCTN1* and *DCTN2* were significantly upregulated in non-small cell lung cancer (NSCLC) and osteosarcoma SJSA-1 cell line, respectively [27,28]. Another study reported that the mRNA expressions of upregulated *DCTN6* and downregulated *DCTN1, DCTN2*, and *DCTN5* were favorable for the prognosis of cutaneous melanoma [29]. High expression of *DCTN4* was also associated with a satisfactory prognosis of colon adenocarcinoma (COAD) [30]. The most common molecular anomalies of LIHC have been reported in the TERT promoter, *TP53, ARID1A, CDKN2A, CTNNB1, AXIN1*, and *CCND1* genes [31,32]. However, there is no extensive analysis of the relationship between the *DCTN* family genes and their oncogenic roles and clinical significance in LIHC.

In this study, we aimed to assess the data available on several public databases to investigate the prognostic and therapeutic perspectives of distinct *DCTN* subunits in LIHC (**Figure 1**). We analyzed the expression pattern, methylation status, and frequency of genetic alterations in *DCTN* genes in LIHC tissues. We also examined the association between *DCTN* expression and LIHC patients’ survival rates and clinical conditions. Moreover, we evaluated the immunophenotypes of *DCTN* genes in the LIHC microenvironment and the functional relevance of the genes co-expressed with *DCTN* genes in LIHC tissues. The experimental findings of this study should guide further laboratory and clinical research on *DCTN*-based diagnostic and therapeutic discoveries in LIHC.

**Figure 1:**
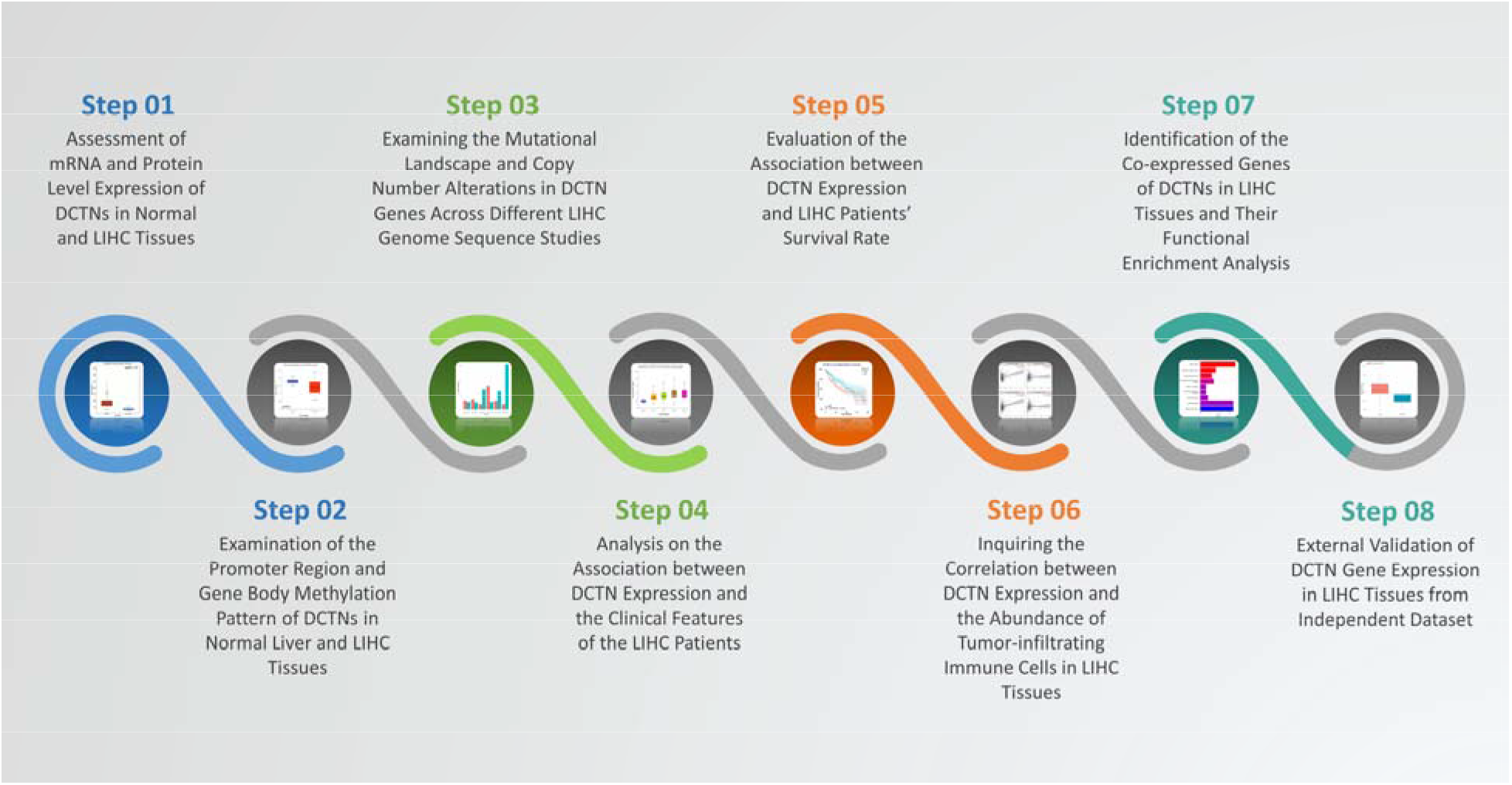
Summary of the strategies employed in the overall study.

## 2. Materials and Methods

### 2.1. Assessment of mRNA and Protein Level Expression of *DCTN*s in Normal and LIHC Tissues

At first, we utilized the OncoDB (http://oncodb.org/) server to retrieve the mRNA level expression pattern *DCTN* in LIHC and adjacent normal liver tissues [33]. The differential expression module of the server was utilized, and the evaluation parameters were kept at default values during the analysis. The OncoDB server collects the RNA sequencing data of respective cancer types from The Cancer Genome Atlas (TCGA) and Genotype-Tissue Expression (GTEx) projects which are then preprocessed, log2 transformed and normalized into transcript per million (TPM) unit. With the aid of the OncoDB tool, we assessed the log2 fold change (log2FC) between the test and control samples upon subjecting them to the student’s t-test. Thereafter, we explored the Expression Atlas server (https://www.ebi.ac.uk/gxa/home) to observe the expression pattern of the *DCTN*s across 22 different LIHC cell lines [34]. The TPM expression values of each respective gene in each cell line were downloaded and then visualized using the ggplot2 package in R studio as a heatmap [35,36]. Finally, the Human Protein Atlas (HPA) (https://www.proteinatlas.org/) web-based tool was utilized to understand the protein level expression pattern of *DCTN* in LIHC and adjacent normal liver tissues [37]. The Pathology and Tissue modules in HPA online tool were inspected to visualize the immunohistochemistry (IHC) images (visualized at 200μm) delineating the protein expression profile of *DCTN*s in the test and control group.

### 2.2. Examination of the Promoter Region and Gene Body Methylation Pattern of *DCTN*s in Normal Liver and LIHC Tissues

We used the University of Alabama at Birmingham Cancer (UALCAN) data analysis Portal UALCAN (http://ualcan.path.uab.edu/) to retrieve the promoter methylation pattern of *DCTN* genes in LIHC and normal lung tissues [38]. The integrated TCGA database was selected for the comparative study, and the hypothesis was subjected to a Student’s *t-*test. Results were obtained in the form of boxplots and then analyzed based on the beta value and p-value cut-off. Next, we explored the University of California Santa Cruz (UCSC) Xena browser (https://xenabrowser.net/) to evaluate the coding sequence expression pattern of *DCTN* genes in LIHC tissues [39]. We selected the TCGA LUAD cohort (n=438) for this analysis. Then, their coding sequence methylation pattern was compared to different LIHC tumor types using the deposited methylation 450k Illumina sequencing data. Finally, we explored the mutation module in Gene Set Cancer Analysis (GSCA) (http://bioinfo.life.hust.edu.cn/GSCA/) to evaluate the correlation between *DCTN* mutation and their expression pattern and LIHC patients’ survival rate [40]. The results were then retrieved as the bubble plot and Kaplan-Meier (KM) plot and analyzed based on the level of significance across the TCGA LIHC dataset.

### 2.3. Examining the Mutational Landscape and Copy Number Alterations in *DCTN* Genes Across Different LIHC Genome Sequence Studies

In this step, we first utilized the cBioPortal web-server (https://www.cbioportal.org/) to understand the frequency of mutations and copy number alterations (CNAs) across the *DCTN* coding regions in LIHC tissues [41]. We selected a total of 6 studies (>1000 samples) involving whole-genome sequencing data on LIHC tissue deposited by MSK, INSERM, TCGA, and others for the analysis of each *DCTN* member. We then explored the OncoPrint and Cancer Type Summary modules to understand the distribution of CNAs and mutations on *DCTN* coding genes across different LUAD studies. We also evaluated the LIHC patients’ overall survival (OS) in relation to *DCTN* mutations. The parameter values were kept default during the analysis in the cBioPortal server. Next, we took the help of the mutation module in GSCA sever to optimize the types of CNAs present across *DCTN* genes in LIHC tissues and any correlation between the present CNAs and their mRNA expression levels. The percent data of each CNA type for *DCTN* genes in TCGA LIHC tissues were downloaded and visualized as a bar diagram with the help of the ggplot2 package in R studio. Later, we retrieved the bubble plot representation of the analysis report on the association between CNAs present in *DCTN* genes and their impact on the mRNA level expression of these genes in LIHC tissues.

### 2.4. Analysis on the Association between *DCTN* Expression and the Clinical Features of the LIHC Patients

We assessed the correlation between *DCTN* expression and LIHC patients’ clinical features again using the UALCAN server. The TCGA database was again used in this experiment to evaluate the impact of *DCTN* expression on the LIHC patients’ clinicopathological features like individual cancer stages, tumor grade, and nodal metastasis status. UALCAN is a web-based resource for studying cancer OMICS data, and it is intended to allow users to find biomarkers or undertake in silico validation of possible genes of interest. The student’s t-test was performed for the formal analysis in this step. The analysis results were retrieved in the form of boxplot and analyzed based on the p-value cut-off.

### 2.5. Evaluation of the Association between *DCTN* Expression and LIHC Patients’ Survival Rate

The Prediction of Clinical Outcome from Genomic Profile (PreCog) (https://precog.stanford.edu/) web-based server was first utilized in this experiment to decipher the correlation between *DCTN* expression and LIHC patients’ OS [42]. The integrated TCGA database for LIHC samples was set at first for the experimental analysis. The results obtained from this server were then visualized in the form of KM plot representation and analyzed based on the hazard ratio (HR) and p-value cut-off. Then, we explored the GEPIA2 web-based server (http://gepia2.cancer-pku.cn/) to understand the association between *DCTN* expression and relapse-free survival (RFS) of LIHC patients [43]. Both the higher and lower threshold exploited as cut-off values were 50% of the mean, and the data were accordingly partitioned into high-expression and low-expression groups. Then we examined the hypothesis by performing a log-rank t-test.

### 2.6. Inquiring the Correlation between *DCTN* Expression in LIHC Tissue and the Abundance of Tumor-infiltrating Immune Cells in LIHC Microenvironment

Here, we utilized the Tumor Immune Estimation Resource 2.0 (TIMER2.0) (http://timer.cistrome.org/) database to assess the correlation between *DCTN* expression and different immune cell infiltration levels in LIHC tissues [44]. TIMER2.0 is a comprehensive resource for systematically analyzing immune infiltrates in various cancer types. The immune module of the server was inspected to establish the correlation between *DCTN* expression and different immune cells, i.e., B cell, CD8+ T cell, CD4+ T cell, Macrophage, Natural Killer (NK) cell in LIHC tissues. The integrated tool called Estimate the Proportion of Immune Cells (EPIC) was considered for the evaluation performed in this step. EPIC collects and correlates the immune cell abundance data from the bulk samples containing a different collection of cells during the gene expression profiling of tumor specimens from the cancer microenvironment [45]. The results of the association study were retrieved in the form of a scatter plot and analyzed based on the Spearmen correlation coefficient and p-value cut-off.

### 2.7. Identification of the Co-expressed Genes of *DCTN*s in LIHC Tissues and Their Functional Enrichment Analysis

In the first step, we identified the top positively co-expressed genes of each *DCTN* family member from the TCGA LIHC dataset (Firehose, Legacy) based on the spearman correlation coefficient from the cBioPortal server. After that, we selected the top 50 positively co-expressed genes of each *DCTN* (total 300) for further analysis. The selected 300 co-expressed genes of all *DCTN*s in LIHC tissues were then subjected to functional enrichment analysis with the help of Clusterprofiler package in R studio delineating their involvement in Biological Processes (BP), Molecular Function (MF), and Cellular Component (CC) inside human body [46]. Later, we also studied the Kyoto Encyclopedia of Genes and Genomes (KEGG) pathway involvement of the overlapping genes.

### 2.8. External Validation of *DCTN* Gene Expression in LIHC Tissues from Independent Dataset

At this step, we testified the expression pattern of *DCTN* genes in two public datasets, i.e., GSE17856 and GSE98383 microarray datasets from the National Center for Biotechnology Information-Gene Expression Omnibus (NCBI-GEO) database. GSE17856 dataset (Platform: Agilent-014850 Whole Human Genome Microarray 4×44K G4112F) contains the total mRNA expression profile from 43 LIHC tissues and 52 adjacent normal liver tissues [47]. On the other hand, GSE98383 (Platform: Affymetrix Human Genome U133 Plus 2.0 Array) comprises the mRNA expression profiles from LIHC patients: 16 LIHC samples, 28 adjacent normal tissue samples, and 30 cirrhosis without LIHC samples [48]. However, only the LIHC and adjacent normal tissue samples were considered during the analysis. The expression data were processed, log2 transformed, normalized and expression values were calculated using the BiocManager package in R studio [49]. Then, the logarithmic fold change between the two variables was calculated using the LIMMA package [50].

## 3. Results

### 3.1. *DCTN*s are Differential Expressed in LIHC Tissues at the mRNA and Protein Levels

The mRNA expression profile of *DCTN*s in normal liver and LIHC tissues was examined from the GEPIA2 server. *DCTN1* was found to overexpress in LIHC tissues (n=372) compared to the adjacent normal tissues (n=50) (log2FC: 0.89, p=1.1e-32) (**Figure 2a**). Similarly, *DCTN2* (log2FC: 1.21, p=6.6e-23), *DCTN3* (log2FC: 0.72, p=7.6e-40), *DCTN4* (log2FC: 0.76, p=1.4e-42), *DCTN5* (log2FC: 0.68, p=1.9e-30) were also found to be overexpressed in LIHC tissues. On the contrary, *DCTN6* was discovered to be under-expressed in LIHC tissues than in adjacent normal liver tissues (log2FC: −0.01, p=6.3e-03). In the next step, we evaluated the expression pattern of all the *DCTN* genes in 22 different LIHC cell lines from the Expression Atlas server. *DCTN2* was observed to be highly expressed in all the selected LIHC cell lines, followed by *DCTN1, DCTN4, DCTN3*, and so forth (**Figure 2b**). Similar to the previous mRNA level observation on *DCTN6*, this gene showed the least expression levels in most of the cell lines. Notably, SNU-182, SNU-387, SNU-398, SNU-423, and SNU-886 were the top observed cell lines in which most of the *DCTN*s showed higher expression. Finally, we investigated the protein level expression of *DCTN*s in LIHC tissues and adjacent normal tissues from the HPA server by inspecting the IHC images prepared from the labeled antibody-*DCTN* protein staining (visualized at 200μm) (**Figure 3**). *DCTN1* exhibited medium staining against the administered antibody in LIHC tissues, whereas no staining was detected in normal tissues. *DCTN2* showed low staining in normal liver tissues and high staining in LIHC tissue. Moreover, the intensity of DCNT4 and *DCTN5* was also lower in normal liver tissues compared to the LIHC tissue. Surprisingly, *DCTN5* showed high IHC staining in both normal liver and LIHC tissue.

**Figure 2:**
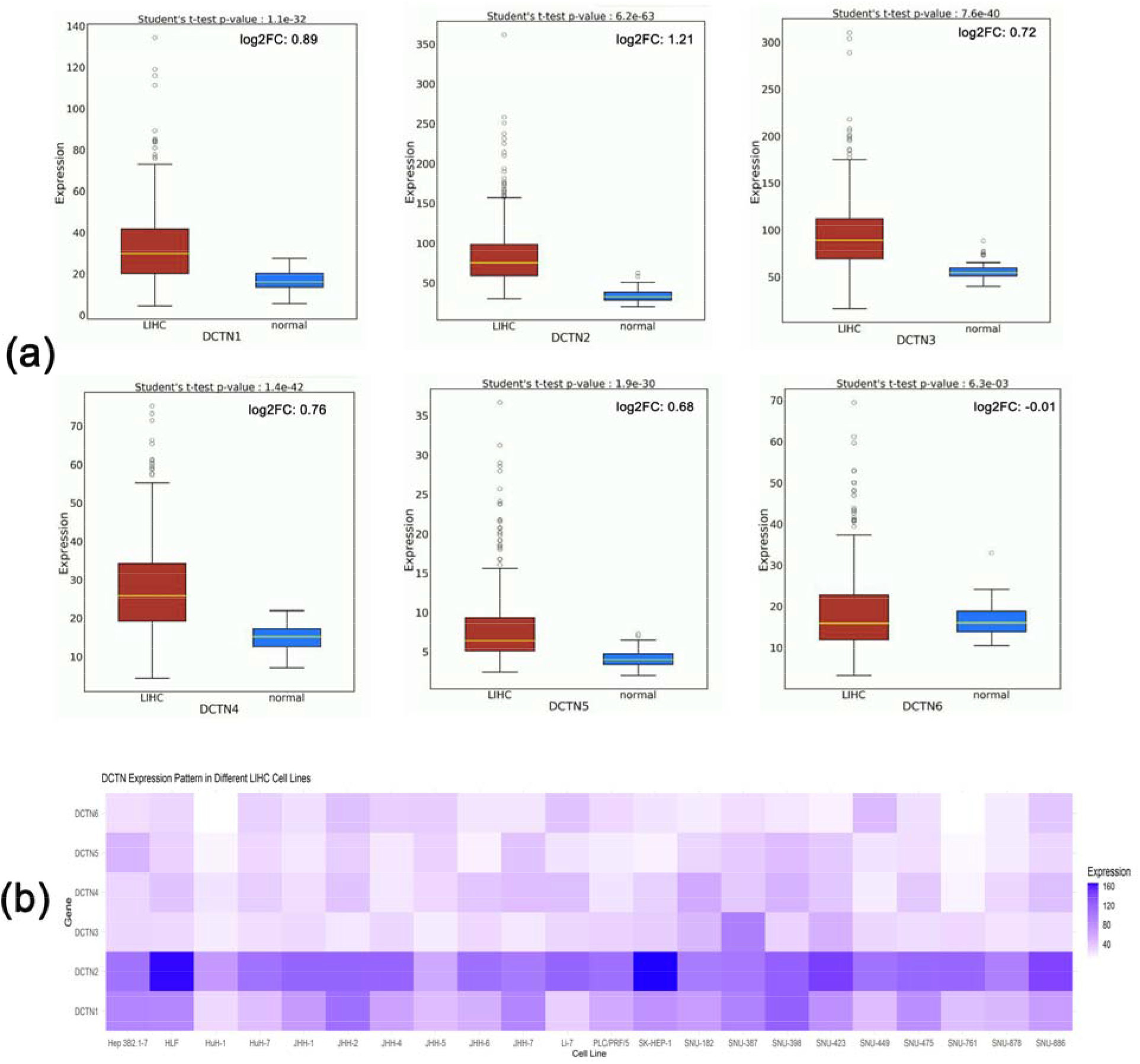
The mRNA level expression pattern of the *DCTN* genes in normal liver (blue) and LIHC (red) tissues (a). The expression values are presented as the log2 normalized transcript per million (TPM) unit. The expression pattern of the *DCTN* genes across different LIHC cell lines (b). The results are presented in the TPM unit. The color gradient represents the expression value of *DCTN*s in TPM units in different cell lines, i.e., low intensity corresponds to a lower TPM, and high intensity corresponds to a higher TPM, while the TPM value escalates with the increasing gradient from low to high.

**Figure 3:**
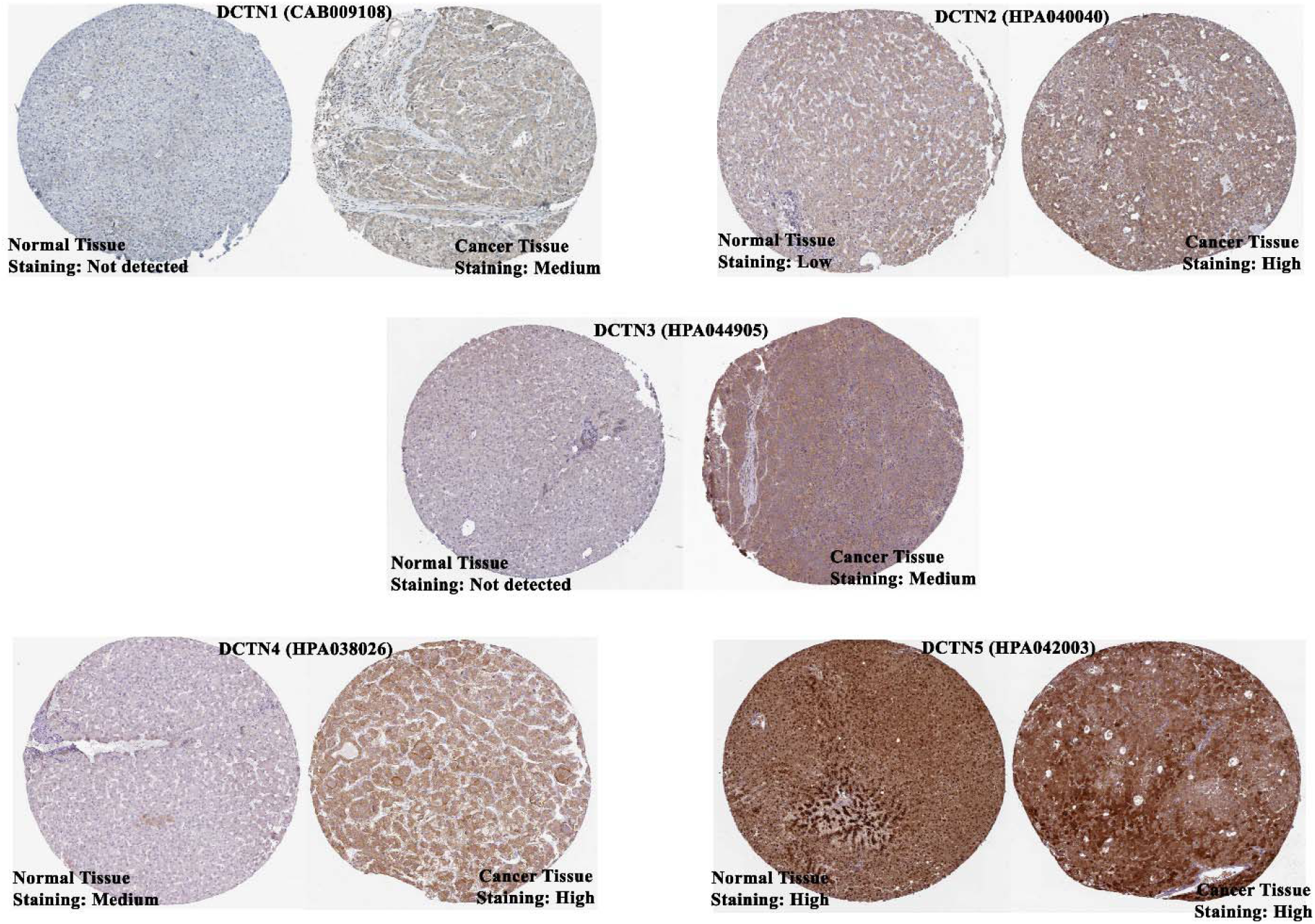
The IHC images of the *DCTN*s in normal liver and adjacent LIHC tissues delineating the protein level expression of *DCTN*s in those samples. The images were visualized at 200 μm in length. The name of the antibody used in the IHC experiment is mentioned within the parentheses beside the gene. The representative IHC image for *DCTN6* was not found in the HPA server.

### 3.2. The Promoters and Coding Sequences of *DCTN*s Have Differential and Distinct Methylation Pattern in LIHC Tissue

*DCTN* coding gene promoter methylation pattern was retrieved from the UALCAN server utilizing the TCGA LIHC cohort. The *DCTN1* coding promoters in LIHC were found to be similarly methylated to those in normal liver tissues though the prediction was not significant (p=1.86e-01) (**Figure 4**). Similarly, no significant observation was also established for the *DCTN3* promoter methylation pattern between the test and control samples, but their promoters were found to be less methylated. On the other hand, the *DCTN2* promoter showed a less methylation pattern in LIHC tissues (n=377) compared to the normal liver tissues (n=50). Moreover, *DCTN4* (p=1e-12), *DCTN5* (p=2.68e-03), and *DCTN6* (p=5.43e-04) promoters all showed less methylation pattern in LIHC tissues compared to the normal liver tissues (**Figure 4**). Thereafter, the gene body of the *DCTN* coding regions was inspected for their methylation pattern from the UCSC Xena browser. *DCTN1* showed that most of the regions of its coding sequence at the 3’ end as indicated by higher beta value marked in red color identifier (**Supplementary Figure S1a**). On the contrary, *DCTN2, DCTN4*, and *DCTN6* indicated that most of the CpG islands in those coding regions might be present sporadically both at the 5’ and 3’ end of the sequences. Interestingly, *DCTN5* contains hypermethylated regions at the 3’ end along the coding sequence (**Supplementary Figure S1a**). The methylation difference analysis on the *DCTN* genes between normal liver and LIHC tissues revealed that *DCTN2* and *DCTN6* gene bodies are less methylated in test samples compared to the control (FDR<0.05) (**Supplementary Figure S1b**). No significant difference between *DCTN1* and *DCTN4* was observed. The level of methylation and mRNA expression correlation analysis reported that unsurprisingly hypomethylation of all the *DCTN* coding genes was positively correlated to the higher mRNA level expression of the genes (**Supplementary Figure S1c)**. Furthermore, the hypomethylation of *DCTN2* (p=0.045) and *DCTN4* (p=0.01) was found to be responsible for the poor progression-free survival of LIHC patients (**Supplementary Figure S1d and S1e)**.

**Figure 4:**
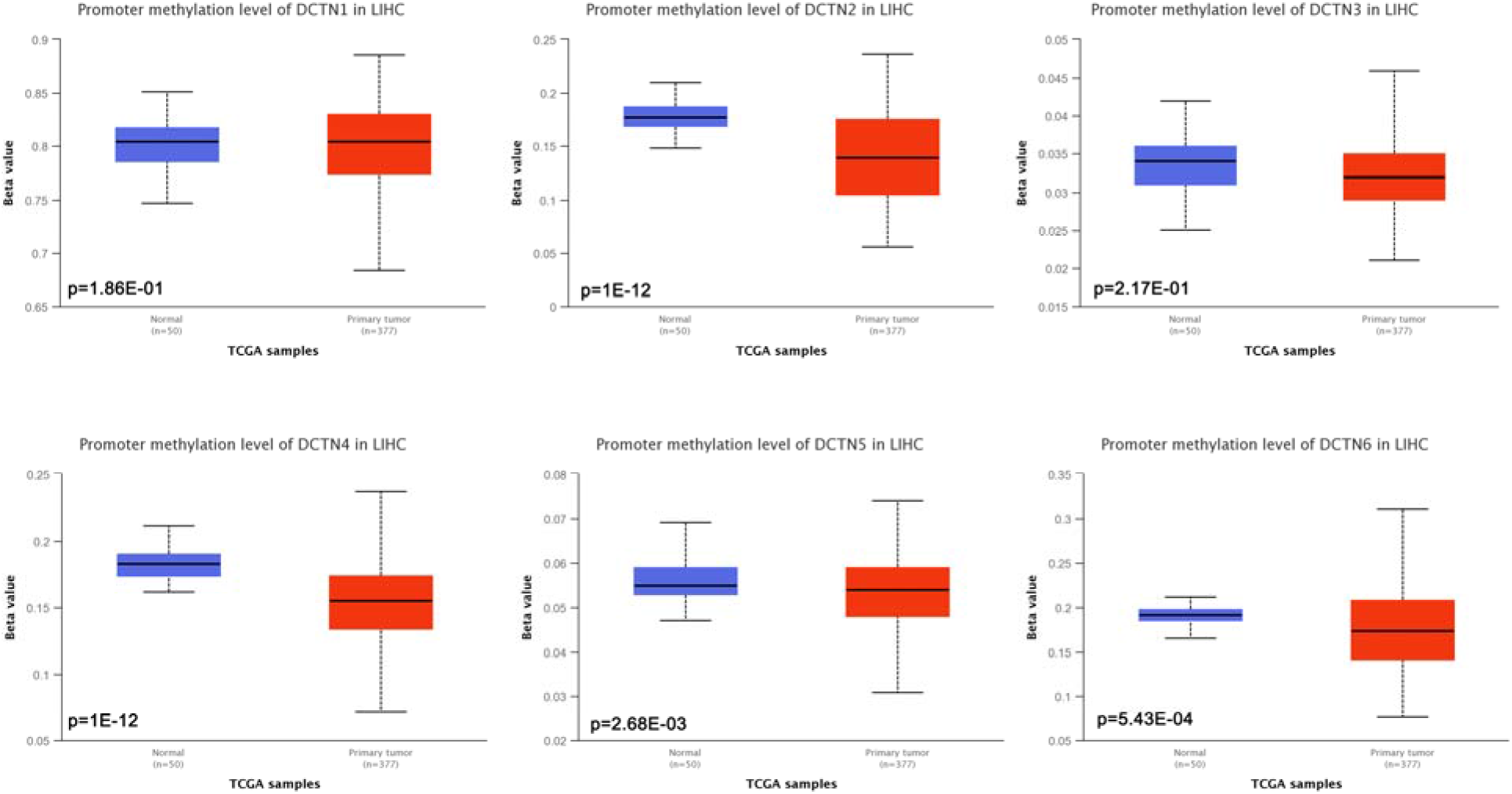
The boxplots representing the differential methylation pattern of *DCTN* gene coding promoters in LIHC tissues compared to the normal tissues. Most of the promoters were found to be less methylated in LIHC tissues. The hypothesis was evaluated by performing a Student’s t-test using the TCGA LIHC cohort through the UALCAN server. Beta value cut-offs in the range of 0.7 to 0.5 indicates hypermethylation, and 0.3–0.25 indicates hypomethylation. Normal: Samples were collected from normal tissues adjacent to the cancerous tissues of LUAD patients within TCGA Cohorts without any demographic and clinical stratification (Source: The Cancer Genome Atlas).

### 3.3. The *DCTN* Coding Genes are Predisposed to Mutation and Copy Number Alteration Events in LIHC Patients

The mutation and copy number alteration (CNA) frequency in *DCTN* genes across different LIHC samples were determined from the cBioPortal server (**Figure 5a**). The analysis report revealed that *DCTN1, DCTN2, DCTN3, DCTN4, DCTN5*, and *DCTN6* had a somatic genetic variation frequency of 1.2%, 1%, 0.4%, 0.4%, 0.2% and 2.5% respectively. Moreover, *DCTN1* and *DCTN2* were predicted to have multiple nonsynonymous mutations within the selected LIHC studies. *DCTN2-5* showed the presence of a devastating form of genetic variation, i.e., amplification (copy number gain). Only *DCTN6*, on the other hand, was predicted to lose the gene coding segment by deep deletion (**Figure 5a**). On the contrary, the analysis report from the GSCA server revealed that all the *DCTN*s underwent both amplification and deletion events in LIHC tissues (**Figure 5b**). Additionally, both the heterozygous amplification and deletion were the most prominent forms of CNAs present in *DCTN* genes compared to homozygous alterations (**Figure 5b**). In summary, *DCTN4* was found to have the most heterozygous amplification events, whereas *DCTN6* showed to undergo the most heterozygous deletion events. Later, we also evaluated the correlation between CNAs present in *DCTN* genes and the LIHC patients’ OS. Though the analysis was not found to be statistically significant, a lower p-value of 0.0535 indicated that the *DCTN* alterations are negatively associated with the OS of LIHC patients (**Figure 5d**). To extend further, the *DCTN* unaltered group of LIHC patients was predicted to have a median OS of ~90 months. whereas that was ~46 months for the altered group of LIHC patients. Finally, the association analysis between the CNAs present in *DCTN* genes and their mRNA expression level unveiled that the presence of CNAs is positively correlated to the mRNA expression levels of the *DCTN* genes in LIHC tissues (FDR<0.05, Cor>0) (**Figure 5e**).

**Figure 5:**
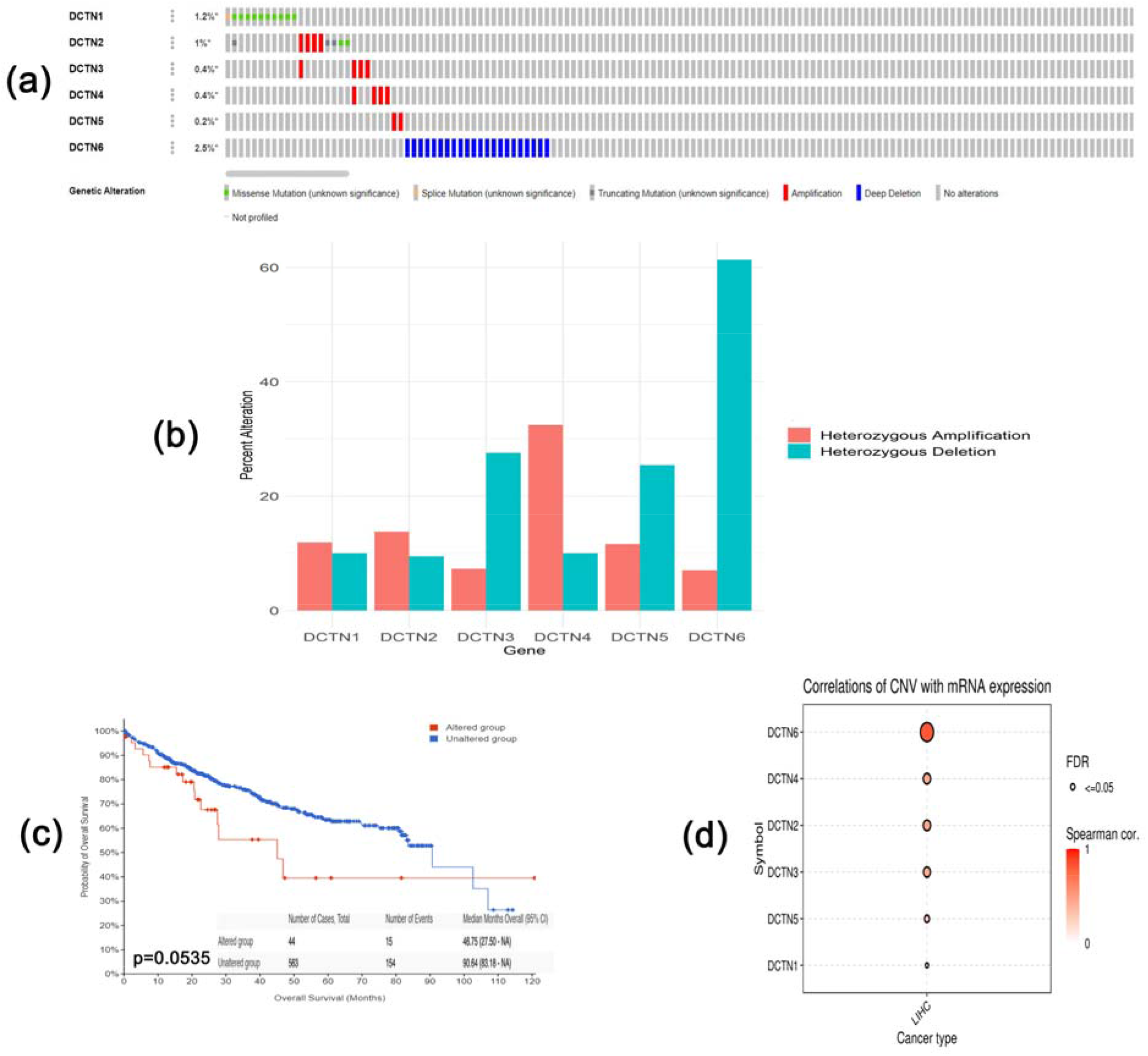
The OncoPrint summary of the frequency of different somatic genetic alterations in *DCTN* genes across different LIHC studies available in the cBioPortal database (a). Bar graph distribution of heterozygous CNAs across *DCTN* genes in LIHC tissues obtained from GSCA server (b). KM plot delineating the association between *DCTN* mutation and LIHC patients’ OS (c). A bubble plot representing the correlation between the CNAs present in *DCTN* genes and their mRNA level expression in LIHC tissues (d).

### 3.4. *DCTN* Overexpression is Associated with Different Clinicopathological Features of LIHC Patients

The association between *DCTN* overexpression and LIHC patients’ clinical features was established using the TCGA LIHC cohort from the UALCAN server. *DCTN1* was found to be overexpressed across all age groups of LIHC patients compared to the normal samples (p<0.01) (**Figure 6a**) (**Supplementary Table S1**). Moreover, *DCTN1* showed a significant rise in the mRNA expression level in accordance with advancing cancer stage (1-4) and LIHC tumor grade (1-4) (p<0.05). *DCTN1* overexpression also showed an association with an advancing metastasis state (N0-N1). However, the association with N1 stage metastasis was not found to be statistically significant (p=5.00E-02) (**Supplementary Table S1**). Similar to *DCTN1, DCTN2*, and *DCTN3* showed overexpression across all age groups of LIHC patients, and the result was predicted to be significant for each clinical parameter (p<0.05) (**Figure 6b** and **6c**). Both *DCTN2* and *DCTN3* showed an increasing level of mRNA expression throughout the advancing cancer stages and tumor grade, although a downward trend in expression level was observed in stage 4 for *DCTN2;* still, the threshold remained above the normal expression value (p<0.05). Overexpression of both the genes was found to be associated with nodal metastasis stages (P<0.05) though the association with the N1 stage for *DCTN2* was not significant (p=9.20E-02) (**Supplementary Table S1**). *DCTN4* and *DCTN5* showed a marginal decrease in mRNA expression from an earlier age (21-40 years group) to the most advanced age group. However, the expression level of those genes was always above those in normal liver tissues (p<0.05). In a similar fashion to other *DCTN* genes, *DCTN4* and *DCTN5* were also reported to be overexpressed and maintained the upward trend in expression with advancing cancer stages and tumor grades (p<0.05) (**Figure 6d** and **6e**). Intriguingly, DCNT6 was found to be declining with the advancing age group though the association with the two most advanced ages was reported to be insignificant (p>0.05) (**Figure 6a**). Moreover, the expression value of *DCTN6* in cancer stages 2 and 4 were predicted to be lower than that of normal liver tissues; again, such association was not found to be significant (p>0.05) (**Supplementary Table S1**). Furthermore, the median expression level of *DCTN6* in grades 1 (p>0.05), 2 (p<0.05), and 4 (p>0.05) of LIHC was lower in comparison with normal expression level, whereas the gene was found to be overexpressed in grade 3 (p<0.05). In terms of nodal metastasis status, *DCTN4-6* showed a significant level of overexpression in the N0 (p<0.001) stage, but no significant association was observed for *DCTN5* and *DCTN6* with expression in the N1 metastasis stage (p>0.05) (**Supplementary Table S1**).

**Figure 6:**
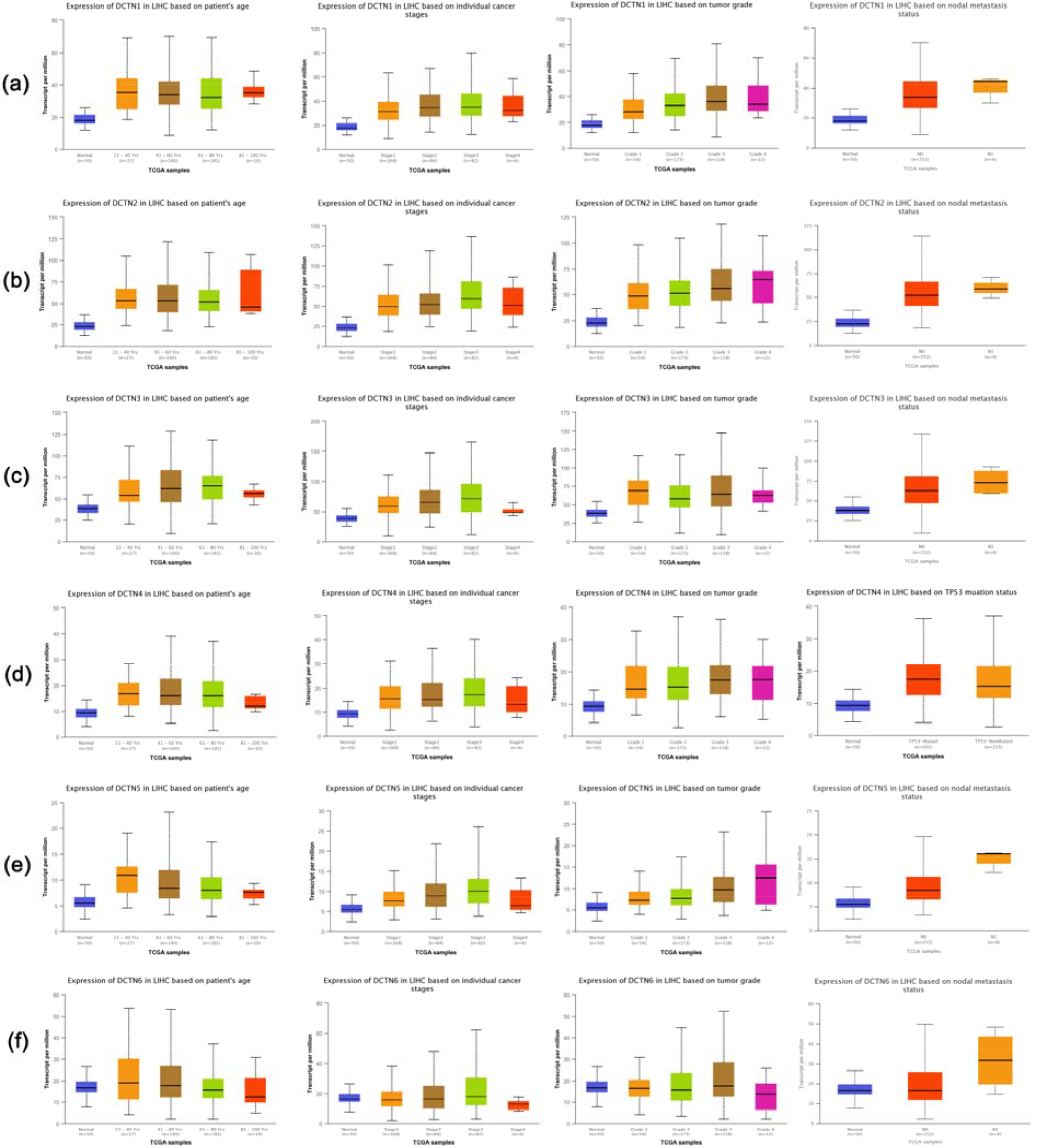
The association between *DCTN* overexpression and different clinical features of LIHC patients. All the *DCTN*s except for *DCTN6* showed noticeable association with most clinical parameters examined. The hypothesis was evaluated by performing a Student’s *t-test* across the TCGA LIHC cohort through the UALCAN server. Normal: Samples collected from normal tissues adjacent to the cancerous tissues of LUAD patients within TCGA Cohorts without any demographic and clinical stratification (Source: The Cancer Genome Atlas).

### 3.5. *DCTN* Expression Can Predict the Overall and Relapse-free Survival of LIHC Patients

The OS analysis on *DCTN1* gene expression in LIHC from the PreCog server revealed that higher gene expression is associated with the unfavorable OS of LIHC patients (HR: 1.32, p=0.044) (**Figure 7**). Additionally, *DCTN1* overexpression was also discovered to be associated with the poor RFS of LIHC patients (HR:1.5, p=0.008) (**Supplementary Figure S2**). In the case of *DCTN2* expression, its overexpression was again found to be associated with the worsening OS of LIHC patients (HR:1.56, p=0.009) (**Figure 7**). Similarly, *DCTN2* overexpression was also reported to be responsible for negative RFS outcomes in LIHC patients (HR:1.5, p=0. 008) (**Supplementary Figure S2**). Similarly, *DCTN3* overexpression was found to be linked to both poor OS (HR: 1.34, p=0.047) and RFS (HR: 1.4, p=0.045) in LIHC patients (**Figure 7**, **Supplementary Figure S2**). However, though the overexpression of *DCTN4* was found to be associated with the poor OS (HR: 1.32) of LIHC patients, the correlation was not observed to be significant (p>0.05) (**Figure 7**). Likewise, the association between *DCTN4* expression and LHC patients’ RFS was also not discovered to be significant, whereas, surprisingly, the analysis indicated that its overexpression may predict better RFS of LIHC patients (HR:0.97, p=0.86) (**Supplementary Figure S2**). In terms of *DCTN5* expression, its overexpression was again observed to be significantly correlated with the poor OS (HR:1.46, p=0.028) and RFS (HR:1.5, p=0.01) of LIHC patients (**Figure 7** and **Supplementary Figure S2**). Though the *DCTN6* expression and its correlation with LIHC patients’ poor OS was not found significant, a very lower p-value of logrank t-test signified that the *DCTN6* overexpression may also predict the OS outcome of LIHC patients (HR: 1.31, p=0.056) (**Figure 7**). Additionally, our analysis revealed that *DCTN6* might predict poor RFS of LIHC patients with a considerable and lower p-value of RFS analysis and HR>1 (HR:1.3, p=0.09) (**Supplementary Figure S2**).

**Figure 7:**
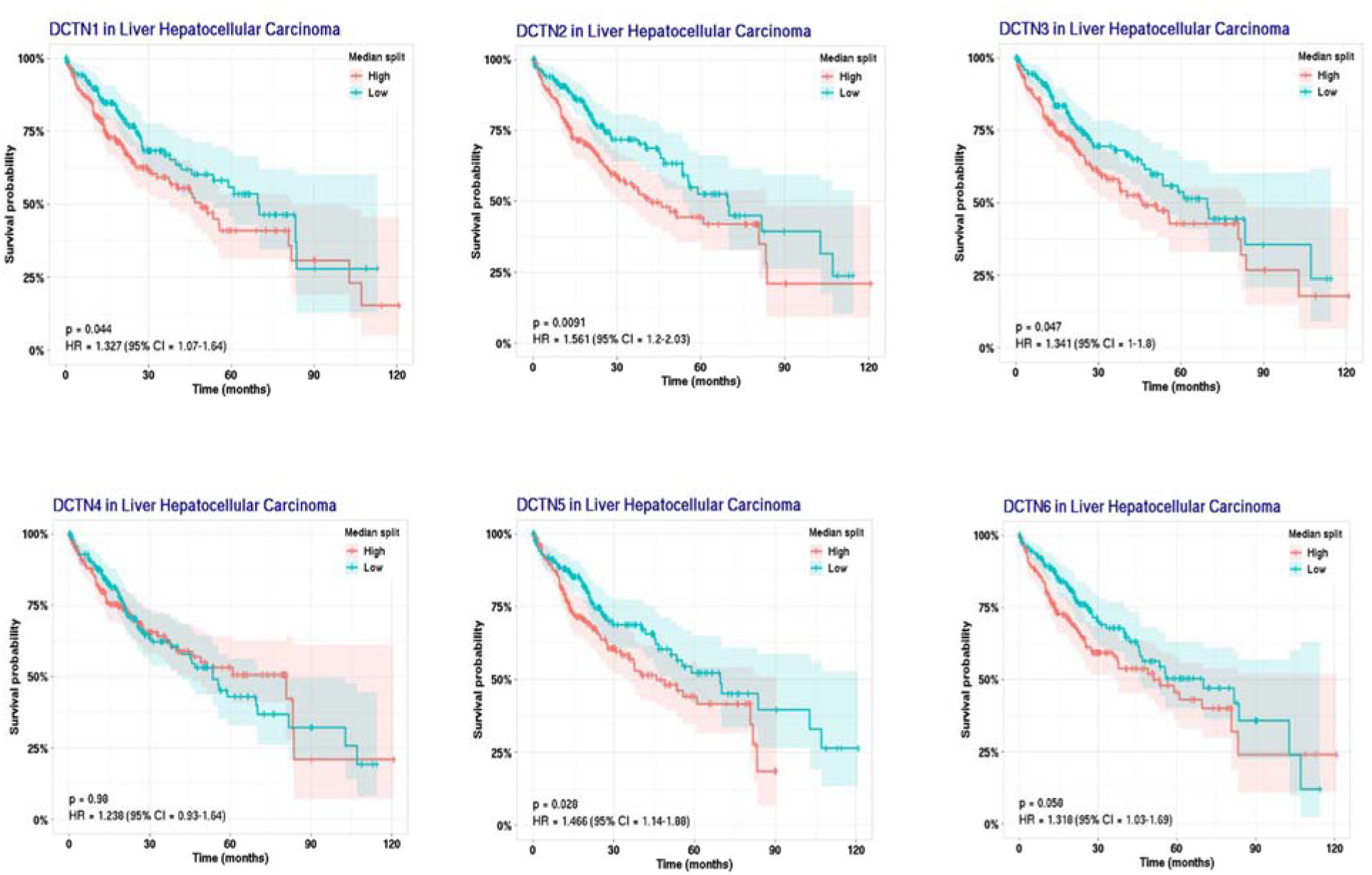
KM plot representation of the association between *DCTN* family gene expression and LIHC patients’ OS. The red and green plot represents the higher and lower *DCTN* expressing LIHC patients, respectively. The vertical mark along the plot indicates an event (death). Overexpression of the *DCTN*s was found to be significantly (except for *DCTN4* and *DCTN6*) associated with the poor OS of LIHC patients.

### 3.6. *DCTN* Expression is Associated with the Abundance of Tumor-infiltrating Immune Cells in LIHC Microenvironment

The association analysis between *DCTN* expression and the infiltration levels of different immune cells in and adjacent tissues of the LIHC microenvironment revealed that the expression of *DCTN1* is significantly and positively correlated with the abundance of B cells (Cor:0.349, p=2.46e-11), CD8+ T (Cor:0.17, p=1.51e-03) cell and CD4+ T cell (Cor:0.154, p=4.09e-03) (**Figure 8a**). On the contrary, *DCTN1* expression in LIHC tissues was observed to be negatively correlated with the abundance level of Macrophages (Cor: −0.434, p=2.60e-17) and NK cells (Cor: −0.112, p=3.77e-02) (**Figure 8a**). *DCTN2* expression was also positively and significantly associated with the infiltration levels of B cell (Cor:0.337, p=1.31e-10), CD8+ T (Cor:0.156, p=1.68e-03) cell and CD4+ T cell (Cor:0.223, p=2.94e-05) in LIHC microenvironment (**Figure 8b**). Similar to *DCTN1* expression, *DCTN2* expression in LIHC tissues was also discovered to be negatively correlated with the infiltration level of Macrophages (Cor: −0.586, p=3.90e-33) and NK cells (Cor: −0.169, p=1.68e-03) (**Figure 8b**). *DCTN3* expression was positively correlated with the abundance level of B cell (Cor:0.241, p=5.78e-06) and CD8+ T (Cor:0.284, p=8.41e-08) and negatively correlated with macrophage (Cor: −0.422, p=2.58e-16) infiltration level (**Figure 8c**). However, the association between the *DCTN3* expression levels and the abundance level of NK cells and CD4+ T cells was not found to be significant (p>0.05) (**Figure 8c**). Alongside, the expression level of *DCTN4* was discovered to be positively associated with B cell, CD8+ T cell, and CD4+ T cell (Cor: >0.130, p<0.05) and negatively associated with macrophage and NK cell infiltration levels (Cor: <-0.240, p<0.01) (**Figure 8d**). The expression of *DCTN5* expression was significantly and positively correlated with the abundance of B cell (Cor:0.414, p=1.05e-15), CD8+ T (Cor:0.15, p=5.27e-03) cell and CD4+ T cell (Cor:0.192, p=3.36e-04) (**Figure 8e**). However, *DCTN5* expression was observed to be negatively correlated with the abundance level of macrophages (Cor: −0.542, p=2.60e-17) and NK cells (Cor: −0.168, p=1.71e-03) (**Figure 8e**). Similar to all other *DCTNs, DCTN6* expression was also found to be significantly and positively associated with B cell, CD8+ T cell, and CD4+ T cell (Cor: >0.178, p<0.01) and negatively associated with macrophage and NK cell infiltration levels (Cor: <-0.164, p<0.05) (**Figure 8f**). Overall, most of the *DCTN*s showed a strong positive correlation with B Cell and CD8+ T cell and a strong negative correlation with macrophage infiltration levels in the LIHC microenvironment.

**Figure 8:**
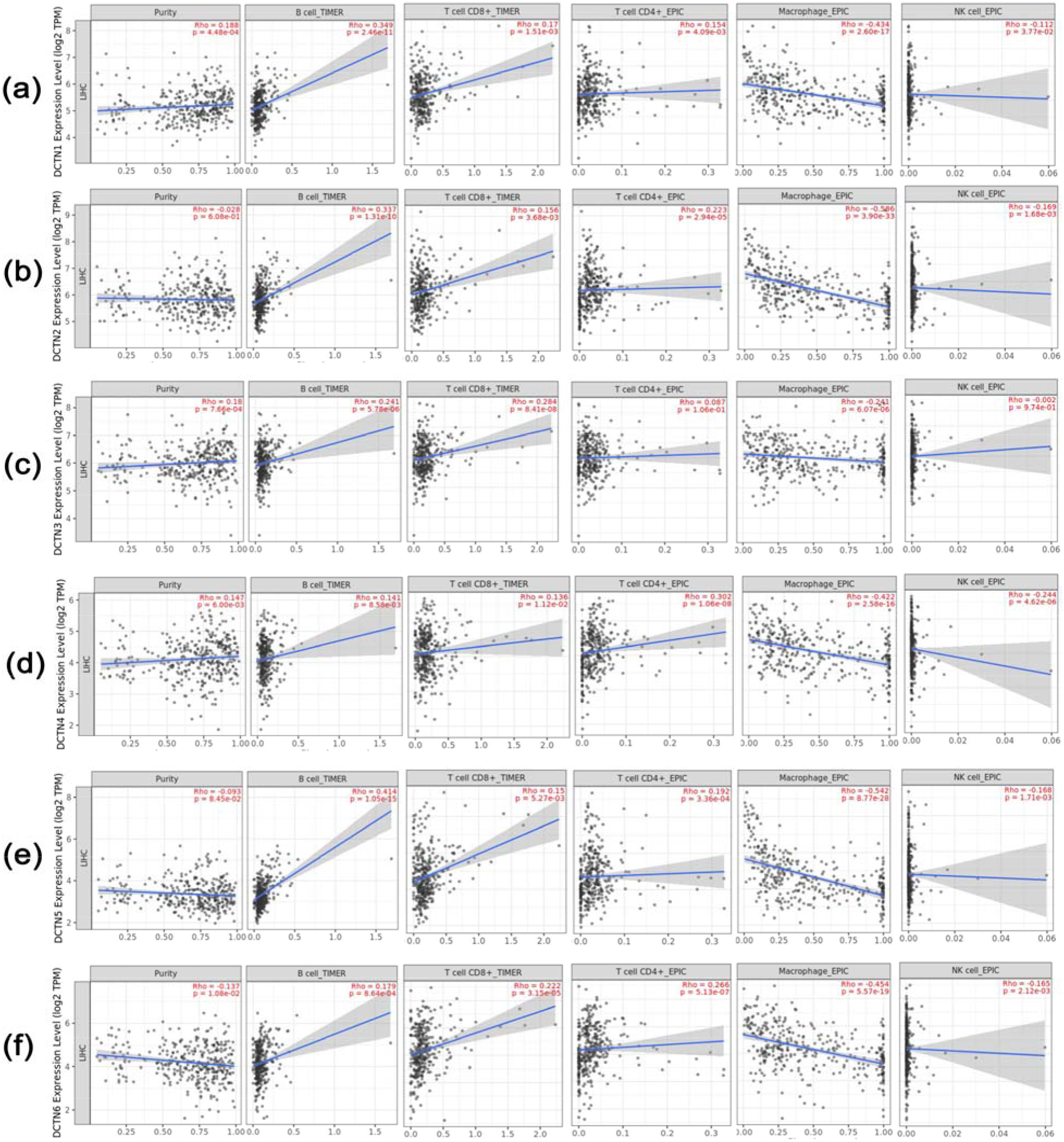
Scatter plot representation of the association between *DCTN* expression and different immune cell infiltration levels in LIHC tissues: (a) *DCTN1*, (b) *DCTN2*, (c) *DCTN3*, (d) DTN4, (e) *DCTN5*, (f) *DCTN6*. Varying degrees of significant positive and negative association were observed between *DCTN* expression levels and different immune cell infiltration levels in LIHC tissues.

### 3.7. The Positively Co-expressed Genes of *DCTN*s in LIHC Tissues and Their Functional Relevance

The analysis of the top co-expressed genes of *DCTN*s in LIHC tissues revealed that *GTF3C2* is the most highly co-expressed gene of *DCTN1* (Cor:0.48, p=6.25e-22) (**Supplementary Figure S3a)**. *DCTN2* and *DCTN3* showed highest co-expression with PRR13 (Cor:0.55, p=8.52e-30) and NDUFB6 (Cor:0.71, p=1.17e-55), respectively, in LIHC tissues (**Supplementary Figure S3b** and **S3c)**. *DCTN4* was found to be most highly co-expressed with *SLU7* (Cor:0.61, p=1.64e-37), and *DCTN5* showed the highest co-expression level with *METTL9* (Cor:0.68, p=7.73e-50) (**Supplementary Figure S3d** and **S3e)**. Additionally, *DCTN6* was reported to be most highly co-expressed with the *PP2R2A* gene (Cor:0.78, p=6.94e-74) in LIHC tissues **(Supplementary Figure S3f)**. Thereafter, the top 50 positively co-expressed genes of each *DCTN* were utilized in the functional enrichment analysis. The KEGG pathway analysis on the co-expressed genes revealed that the cluster was most significantly involved in maintaining tight junction, lysine degradation, pyrimidine metabolism, glycolysis/gluconeogenesis, activities of peroxisome, RNA degradation, and so forth. Moreover, a notable number of genes were also found to be involved in the progression of neurodegenerative diseases, i.e., Huntington’s disease (**Figure 9a**). The biological processes (BP) analysis of the top co-expressed genes reported that the genes are predominantly involved in producing metabolites and energy, purine and pyrimidine metabolic processes, regulation of chromosome organization, ribose phosphate metabolic processes, pyruvate metabolic processes, telomere maintenance, and many other crucial biological functions (**Figure 9b**). Moreover, histone modification activity, enzymatic activity, i.e., transferase activity, damaged DNA binding, and NADP binding were reported to be the major molecular functions (MF) of the cluster of co-expressed genes of *DCTN*s in LIHC tissues (**Figure 9c**). Additionally, the co-expressed genes were mostly found to be operating in the nuclear speck, organelle envelope lumen, tertiary granule lumen, and mitochondrial intramembrane space as observed from the cellular component analysis on the top co-expressed genes of *DCTN*s in LIHC tissues (**Figure 9c**).

**Figure 9:**
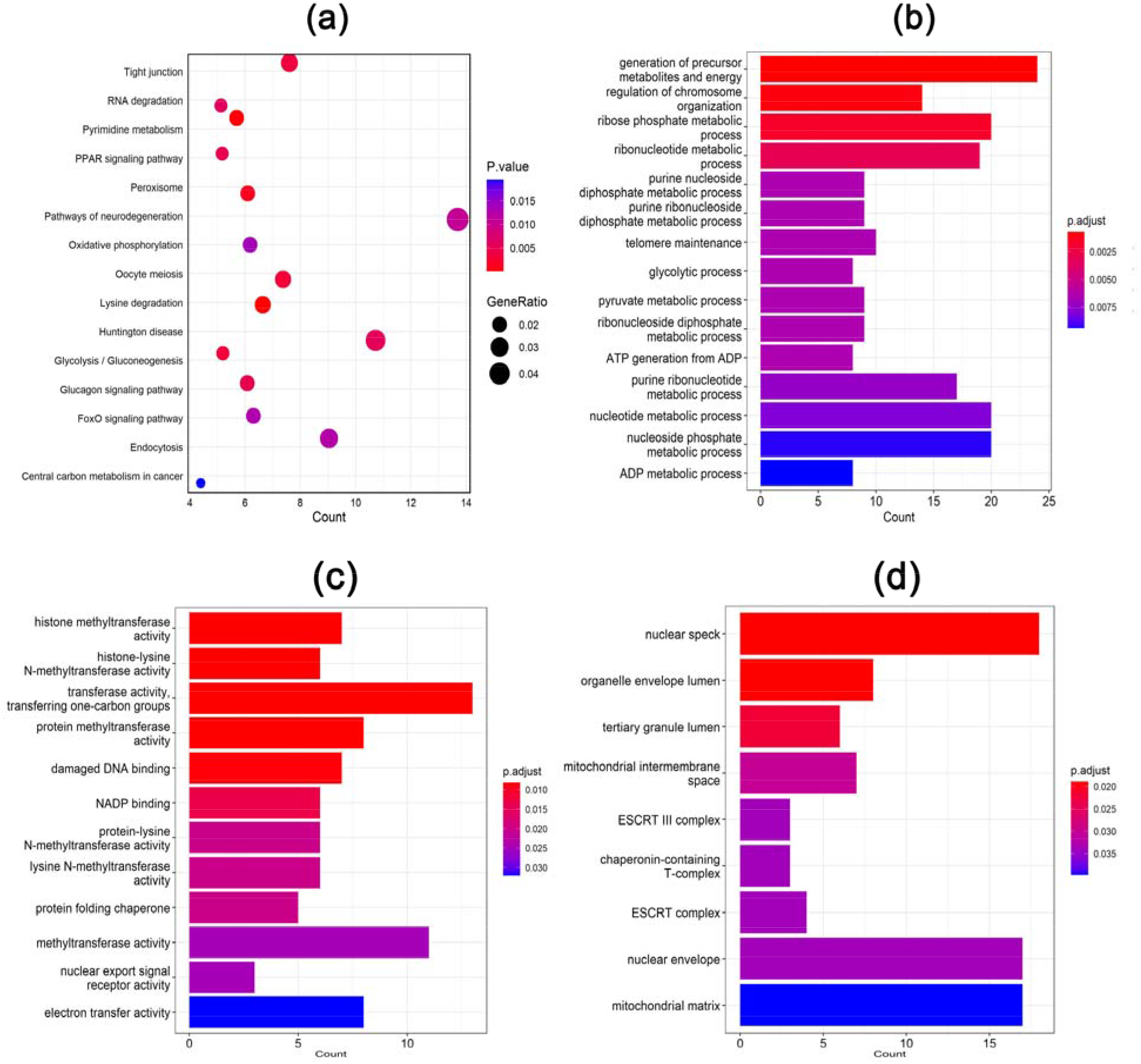
(a) Dot plot representation of the result of KEGG pathway analysis on the top positively co-expressed genes of *DCTN*s in LIHC tissues. Barplot representation of the functional enrichment analysis, i.e., Biological Processes (b), Molecular Function (c), Cellular Component (d) on the neighbor genes of *DCTN*s in LIHC tissues.

### 3.8. Validation of the *DCTN* Expression Pattern in Independent LIHC Datasets

The evaluation of *DCTN*s expression in GSE17856 microarray dataset indicated that *DCTN1* is significantly overexpressed in LIHC tissues (n=43) compared to the adjacent normal tissues (n=52) (log2FC: 0.99, p=0.025) (**Figure 10a**). Similarly, *DCTN2* (log2FC: 0.93, p<0.001), *DCTN3* (log2FC: 0.97, p=0.03), *DCTN4* (log2FC: 0.88, p<0.001) and *DCTN5* (log2FC: 0.97, p=0.01) were also found to be significantly overexpressed in LIHC tissues than in adjacent normal liver tissues (**Figure 10**). However, *DCTN6*, as in par with the mainstream analysis, was also reported to be under-expressed in LIHC tissues (log2FC: −0.21, p=0.02). The analysis of *DCTN* expression in GSE98383 dataset also revealed that *DCTN1* (log2FC: 0.92, p=0.01) and *DCTN2* (log2FC: 0.88, p=0.02) are highly expressed in LIHC tissues (n=16) compared to the adjacent normal tissues (n=28) (**Supplementary Figure S4**). In case of *DCTN3*, though it was found to be overexpressed in LIHC tissues, the association was not discovered to be significant (log2FC: 0.98, p=0.09). However, both *DCTN4* (log2FC: 0.81, p=0.008) and *DCTN5* (log2FC: 0.89, p=0.015) were reported to be overexpressed in LIHC tissues than in normal tissues (**Supplementary Figure S4**). On the contrary, *DCTN6* was again found to be downregulated in LIHC tissues compared to the adjacent normal liver tissues (log2FC: −0.69, p=0.005).

**Figure 10:**
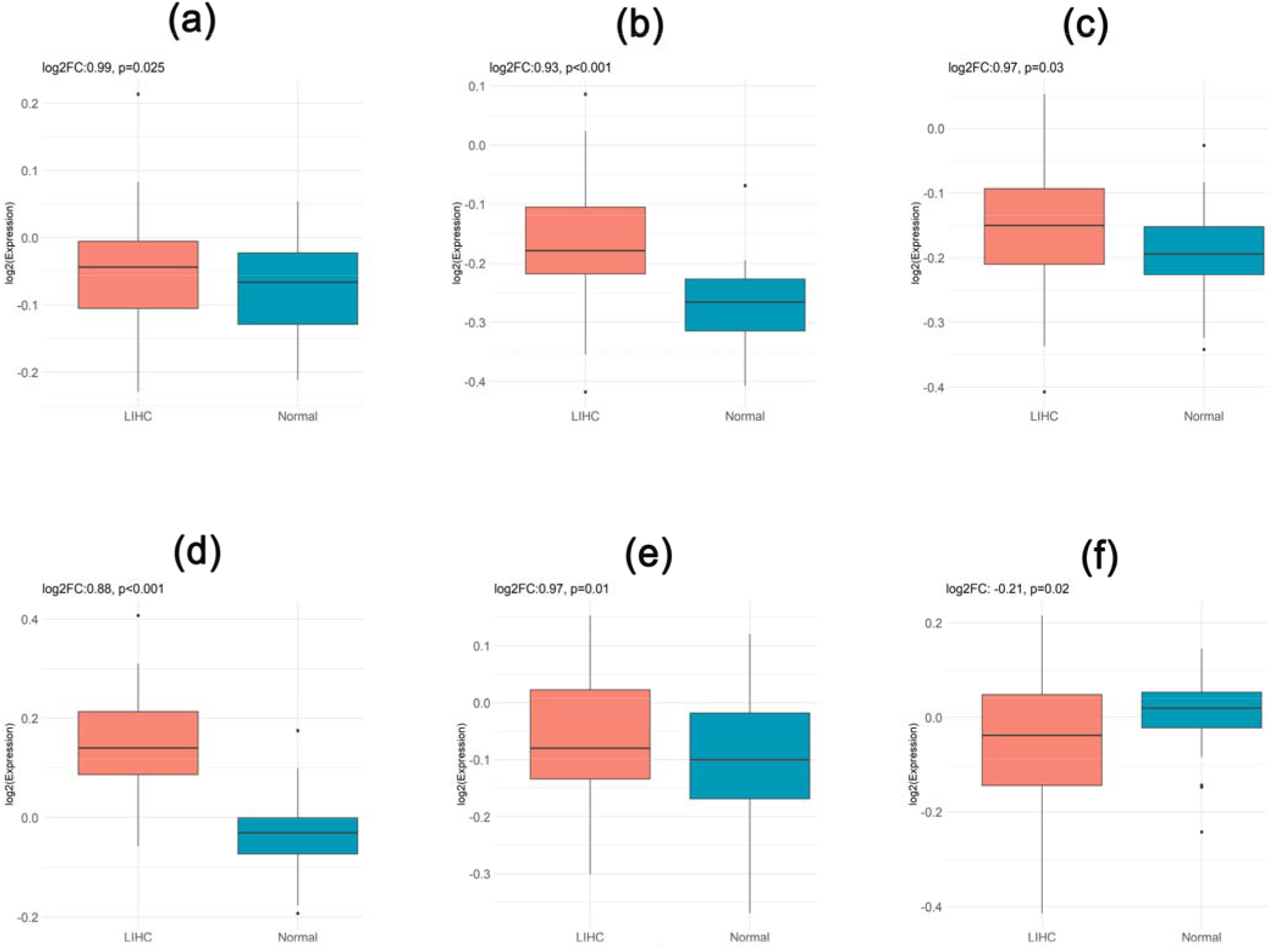
The expression pattern of *DCTN*s in LIHC tissues (n=43) and adjacent normal liver tissues (n=52) from GSE17856 microarray dataset: (a) *DCTN1*, (b) *DCTN2*, (c) *DCTN3*, (d) *DCTN4*, (e) *DCTN5*, (f) *DCTN6*. All the genes except *DCTN6* were found to be significantly overexpressed in LIHC tissues.

## 4. Discussion

This study employed a multi-omics database mining pipeline to explore the possible oncogenic roles and prognostic relevance of the *DCTN* family of genes in LIHC. Differential gene expression analysis at the mRNA and protein level can assist in understanding the potential roles of different genes in multi-factorial, heterogeneous, and complex diseases like cancer [51, 52]. Again, due to the heterogenic nature of cancer and multistep processes involved in the oncogenic development of healthy cells, identifying differentially expressed genes in cancer tissues and their subsequent analysis in relation to patient’s survival can help optimize the underlying mechanism involved in cancer development and progression [51]. In this study, all the selected *DCTN*s were found to be differentially expressed (*DCTN1-5*: upregulated; *DCTN6*: downregulated) in LIHC tissues compared to the adjacent normal tissues at both mRNA and protein levels, suggesting the possible oncogenic roles of *DCTN*s in LIHC (**Figure 2** and **3**). Most of the selected *DCTN*s except *DCTN6* were also observed to be overexpressed in different LIHC cell lines. From the diagnostic perspective, the differential level of *DCTN* mRNA expression observed in this study holds great promise in capturing the disease state of LIHC patients. Additionally, the differential protein level expression of *DCTN*s in LIHC tissues as witnessed by investigating the IHC images from HPA server suggests that *DCTN*s may assist in the proper stratification of LIHC patients, detection of the origin of LIHC metastasis, and tracking the response to any future *DCTN-based* therapeutics against LIHC (**Figure 3**) [53].

Aberrant DNA methylation is the most studied epigenetic phenomenon in cancer diagnostic and therapeutic development. Promoter and gene body methylation controls the gene activity in different manners, and usually, promoter hypomethylation of a specific gene is associated with increased activated transcription of the gene in question [54, 55]. Additionally, aberrant gene body methylation in cancer cells can also influence the dosage at which a particular gene is transcribed inside healthy cells by affecting the indigenous nucleosome structures within the chromatin architecture [56]. In this study, most of the *DCTN* coding promoters, except for *DCTN1*, were found less methylated (**Figure 4**). Moreover, *DCTN2* and *DCTN6* coding sequences were also observed to be significantly less methylated in LIHC tissues compared to the adjacent normal liver tissues (**Supplementary Figure S1**). Thus, the aberrant methylation patterns of the *DCTN* genes might be accounted for their differential expression levels in LIHC tissues. In support of such a hypothesis, a less methylation pattern of *DCTN* genes was later found to be significantly associated with their mRNA level overexpression in LIHC tissues (**Supplementary Figure S1**). In terms of methylation-sensitive diagnosis, specific CpG sites in different genes are powerful diagnostic signatures in LIHC, and the aberrant methylation pattern is usually observed at an earlier stage of cancer development, while the pathologic testing may not identify the tumorigenic transformation of healthy cells at that stage [57, 58]. Moreover, the abnormal methylation pattern of specific genes can be identified from the bodily fluid of LIHC patients, i.e., serum, and thus this kind of epigenetic marker can help in the non-invasive diagnosis of LIHC patients [59]. In our study, all the *DCTN*s were observed to contain distinct CpG islands across their coding sequence, which, along with their aberrantly methylated promoters, could help in the high-throughput diagnosis of LIHC patients in a non-invasive manner and at an earlier stage (**Supplementary Figure S1**). Alongside, due to the reversible nature of DNA methylation, unlike mutations, methylation-targeted therapy remains an important and promising therapeutic option in LIHC treatment with applied strategies like regulation of DNA methyltransferase activities [60, 61]. In this study, *DCTN2* and *DCTN4* hypomethylation was negatively associated with the poor PFS of LIHC patients suggesting that dysregulated methylation of *DCTN*s could be targeted in the methylation-based treatment of LIHC patients and making the patient-specific epigenetic clinical decision (**Supplementary Figure S1**).

Somatic mutations are one of the major hallmarks in the development and progression of cancer. Different forms of devastating mutations, i.e., amplification, deletion, or splice, collectively known as CNAs, can significantly propel the oncogenic transformation of healthy cells to a greater extent than other point mutations like nonsynonymous mutations [62, 63]. Such CNAs remarkably alter the optimum transcriptional and translational dosage of particular genes and can influence the cancer development processes by alternating the cellular homeostasis [64]. In this study, we also recorded multiple numbers of damaging CNAs, i.e., heterozygous amplification and deletion, along with several point mutations across the coding sequences of most of the *DCTN*s in LIHC tissues that might be responsible for their differential level of expression (**Figure 5**). Though the impacts of *DCTN* mutations on cancer initiation and progression remain largely unknown to date, a recent study has proposed that the *DCTN*s are highly susceptible to mutations in cancer cells [65]. Again, our analyses suggested that *DCTN* mutations are negatively associated with the OS of LIHC patients indicating that the genetic alterations in *DCTN* genes may influence the development and subsequent growth of LIHC while justifying their potential as the therapeutic targets for LIHC. Additionally, the high frequency of different gene mutations, i.e., TERT (alteration frequency: 60%) and CTNNB1 (alteration frequency: ~40%), can predict the LIHC outcome and serve the high-throughput diagnostic perspective of LIHC patients [66]. Hereby, the observed alterations in *DCTN* genes in this study, specifically *DCTN6*, which was reported to undergo high deep deletion events, may assist as a surrogate diagnostic marker in the dual diagnosis of LIHC patients, along with *DCTN*-based other conventional pathologic testing (**Figure 5**).

Later, the survival analysis of the LIHC patients in relation to *DCTN* expression revealed that high expression of *DCTN* genes could predict the poor OS of LIHC patients (**Figure 7**). Alongside, higher expression of most of the *DCTN* genes (except *DCTN4*) was also observed to predict the poor RFS of LIHC patients (**Supplementary Figure S2)**. These pieces of evidence suggest that the *DCTN* gene expression pattern might be adopted to capture the high-risk group of LIHC patients, recommend necessary follow-ups, and measure the therapeutic efficacy of any intervention in LIHC patients. Association analysis between *DCTN* gene expression and LIHC patients’ clinical features revealed that most of the *DCTN*s are highly expressed earlier in LIHC patients. Moreover, a significant association was also observed between *DCTN* overexpression and all different LIHC stages and grades (**Figure 6**). These findings signify that the *DCTN* expression pattern may assist in predicting the cellular fitness of LIHC cells and tracking the LIHC patients throughout the entire clinical course. The correlation between *DCTN* overexpression and LIHC patients’ metastatic states suggests that the expression level of *DCTN*s might also predict the metastatic condition of LIHC.

The association analysis between *DCTN* expression and different immune cell infiltration levels revealed that most of the *DCTN*s were positively and negatively associated with the abundance level of different immune cells in the LIHC microenvironment. More specifically, B and T cells were found to show a significant positive correlation with most of the *DCTN* expression levels in LIHC tissues. A previous study involving 112 LIHC samples reported that the abundance of B and T cells in LIHC patients can significantly improve patients’ prognoses, and a higher titer of those immune cells can predict better OS [67]. Thus, the abundance of different immune cells and their association with *DCTN* expression could further assist in *DCTN*-based LIHC diagnosis or immunotherapy decisions such as adoptive T cell transfer or immune checkpoint inhibition by modulating the activity of T cells [68]. On the contrary, the mutated version of *DCTN*s in LIHC patients may alter the normal immune activity and thus can affect the prognosis of LIHC patients.

The analysis of the positively co-expressed genes of *DCTN*s revealed that GTF3C2 is the top co-expressed gene of *DCTN1* in LIHC tissues (**Supplementary Figure S3**). Though the absolute oncogenic roles of this gene remain unstudied to date, a recent systematic study has proposed that GTF3C2 may be involved in colorectal cancer (CRC) development, and its overexpression could predict better OS of CRC patients [69]. A recent study with gastric cancer samples (n=103) reported that the expression of NDUFB6, a gene most positively co-expressed with *DCTN3* in LIHC tissues, is associated with the invasion and distant metastasis of gastric cancer patients [70]. SLU7, a gene found to be highly co-expressed in LIHC tissues with *DCTN4*, is essential to maintaining liver cell homeostasis, and its overexpression has been linked to promoting proliferation, migration, epithelial-mesenchymal transition, and reducing apoptosis rate in LIHC cells in studies involving clinical specimens [71,72]. The top positively co-expressed gene of *DCTN5* in LIHC tissues, METTL9, promotes cancer cell growth by methylating histidine residues and its deletion reduces the proliferation of tumor cells [73]. Moreover, the PPP2R2A gene was highly co-expressed with *DCTN6* in LIHC tissues. The down expression and mutation of this gene are associated with colorectal, pancreatic, and breast cancer exacerbation and poor OS in breast cancer patients [74–76]. Hereby, given that the co-expressed genes are functionally related, we hypothesize that differential *DCTN* expression may also affect the LIHC progression by altering the expression levels of their neighbor genes similarly as observed in other cancer types. Afterward, the functional enrichment analysis of the top co-expressed genes of *DCTN*s in LIHC tissues revealed that most of the genes were predominantly involved in crucial biological processes like nucleotide, i.e., purine and pyrimidine metabolism, damaged DNA binding, chromosome organization, histone methyltransferase activity, glycolytic processes, maintaining tight junction and peroxisome activity and so forth (**Figure 9**). The deregulation of such activities could also lead to the oncogenic transformation of healthy cells [77–80]. Therefore, the co-expressed genes of *DCTN*s in LIHC tissues could also be investigated while extending clinical development procedures on making *DCTN-based* diagnostic and therapeutic measures for LIHC. Lastly, the *DCTN* expression pattern was validated in two small-scale (n=~20-50) public microarray datasets of LIHC studies, i.e., GSE17856, and GSE98383, whereas the mainstream analysis was projected on a large-scale (n=~400-600) RNA-sequencing data from TCGA and GTEx cohorts. The evaluation step also indicated that the expression pattern of the *DCTN*s (except *DCTN5* in GSE98383) in the independent dataset was also at par with our preliminary experimental findings (**Figure 10**).

Overall, this study demonstrated that *DCTN*s are differentially expressed (*DCTN1-5:* upregulated, *DCTN6:* down-regulated) at both mRNA and protein levels in LIHC tissues compared to the adjacent normal tissues. Most of the *DCTN*s were also reported to overexpress in different cell lines. Additionally, *DCTN*s were recorded to be overexpressed in LIHC patients at an earlier stage across different LIHC stages and Grades. Overexpression of *DCTN*s was mostly associated with poor OS and RFS of LIHC patients. Therefore, it is quite reasonable to postulate that the transcriptomic and proteomic differential level of *DCTN* expression may assist in the *DCTN-based* diagnosis of LIHC patients. Furthermore, since *DCTN*s are involved in crucial cell cycle processes [23–26], their abnormal expression pattern might affect the optimum controls of cell division and progresses the healthy cells through the oncogenic transformation which signifies their potential as the targets for therapeutic interventions in LIHC patients. Alongside, the promoters and coding sequences of *DCTN*s were observed to be differentially methylated and mutated in LIHC tissues. Such genetic and epigenetic alterations in *DCTN* genes should further assist in increasing the precision of *DCTN*-based LIHC diagnosis and patient-specific clinical decisions. The immunophenotype analysis of *DCTN* genes in the LIHC microenvironment revealed their association with a variety of immune cell infiltration levels. And functionally co-expressed genes of *DCTN*s were discovered to be involved in oncogenic processes in different cancer types and maintaining biological processes, which may lead to cancer development upon deregulation. Thereby, the immunophenotypes and functionally related genes of *DCTN*s in LIHC tissues should also be investigated since these signatures could guide the formulation of any possible dual diagnosis or combinatorial therapeutic approaches against LIHC. Finally, this study recommends that *DCTN*s and their transcriptional and translational products are potential candidates for LIHC diagnosis and treatments. The scientific findings of the present study will help further the clinical development of *DCTN-based* diagnostic and therapeutic measure discovery for LIHC.

Lastly, our study involved a large number of omics data to explore the therapeutic and prognostic potentials of *DCTN*s in LIHC for the first time. Most of the analysis was found to be significant in between test and control variables. Moreover, this study provided a multi-omics (i.e., genomic, transcriptomic, proteomic) overview of *DCTN* gene expression and its clinical relevance in LIHC. However, our study has some limitations, i.e., it cannot provide a clearer picture of the specific molecular pathogenesis of *DCTN* gene expression in LIHC. Additionally, it cannot confer the degrees at which the genetic or epigenetic alterations in *DCTN* genes may affect their expression levels in LIHC tissues that require further laboratory inspection which is presently operative by the authors.

## Authors’ Contributions

Conceptualization and experimental designing: M.A.U. Experiment, analysis and interpretation: M.A.U., T.T., N.N.I and B.K. Writing, review and editing: M.A.U., T.T., A.R., N.N.I., M.N.P., A.T.M., and B.K. Supervision and funding acquisition: B.K. All authors have read and agreed to the published version of the manuscript.

## Ethics Approval and Consent to Participate

Not Applicable

## Consent for Publication

Not Applicable

## Availability of Data and Material

All the data are provided within the manuscript and the supplementary material.

## References

1. Bray F, Ferlay J, Soerjomataram I, Siegel R, Torre L, Jemal A. Global cancer statistics 2018: GLOBOCAN estimates of incidence and mortality worldwide for 36 cancers in 185 countries. CA: A Cancer Journal for Clinicians. 2018;68(6):394–424.

2. Chuang S, Vecchia C, Boffetta P. Liver cancer: Descriptive epidemiology and risk factors other than HBV and HCV infection. Cancer Letters. 2009;286(1):9–14.

3. Singh A, Kumar R, Pandey A. Hepatocellular Carcinoma: Causes, Mechanism of Progression and Biomarkers. Current Chemical Genomics and Translational Medicine. 2018;12(1):9–26.

4. Lauby-Secretan B, Scoccianti C, Loomis D, Grosse Y, Bianchini F, Straif K. Body Fatness and Cancer — Viewpoint of the IARC Working Group. New England Journal of Medicine. 2016;375(8):794–798.

5. Ferlay J, Shin H, Bray F, Forman D, Mathers C, Parkin D. Estimates of worldwide burden of cancer in 2008: GLOBOCAN 2008. International Journal of Cancer. 2010;127(12):2893–2917.

6. Bertuccio P, Turati F, Carioli G, Rodriguez T, La Vecchia C, Malvezzi M et al. Global trends and predictions in hepatocellular carcinoma mortality. Journal of Hepatology. 2017;67(2):302–309.

7. Liu Z, Jiang Y, Yuan H, Fang Q, Cai N, Suo C et al. The trends in incidence of primary liver cancer caused by specific etiologies: Results from the Global Burden of Disease Study 2016 and implications for liver cancer prevention. Journal of Hepatology. 2019;70(4):674–683.

8. Akinyemiju T, Abera S, Ahmed M, Alam N, Alemayohu M, Allen C et al. The Burden of Primary Liver Cancer and Underlying Etiologies From 1990 to 2015 at the Global, Regional, and National Level. JAMA Oncology. 2017;3(12):1683.

9. de Martel C, Maucort□Boulch D, Plummer M, Franceschi S. World wide relative contribution of hepatitis B and C viruses in hepatocellular carcinoma. Hepatology. 2015;62(4):1190–1200.

10. Kew M. Hepatocellular carcinoma in African Blacks: Recent progress in etiology and pathogenesis. World Journal of Hepatology. 2010;2(2):65.

11. Forner A, Llovet J, Bruix J. Hepatocellular carcinoma. The Lancet. 2012;379(9822):1245–1255.

12. Palmer D. Radiofrequency Ablation With or Without Transcatheter Arterial Chemoembolization. Journal of Clinical Oncology. 2013;31(21):2756–2756.

13. Jackson R, Psarelli E, Berhane S, Khan H, Johnson P. Impact of Viral Status on Survival in Patients Receiving Sorafenib for Advanced Hepatocellular Cancer: A Meta-Analysis of Randomized Phase III Trials. Journal of Clinical Oncology. 2017;35(6):622–628.

14. Bouattour M, Mehta N, He A, Cohen E, Nault J. Systemic Treatment for Advanced Hepatocellular Carcinoma. Liver Cancer. 2019;8(5):341–358.

15. Balogh J, Victor D, Asham E, Burroughs S, Boktour M, Saharia A et al. Hepatocellular carcinoma: a review. Journal of Hepatocellular Carcinoma. 2016;Volume 3:41–53.

16. Gill S, Schroer T, Szilak I, Steuer E, Sheetz M, Cleveland D. Dynactin, a conserved, ubiquitously expressed component of an activator of vesicle motility mediated by cytoplasmic dynein. Journal of Cell Biology. 1991;115(6):1639–1650.

17. Eckley D, Gill S, Melkonian K, Bingham J, Goodson H, Heuser J et al. Analysis of Dynactin Subcomplexes Reveals a Novel Actin-Related Protein Associated with the Arp1 Minifilament Pointed End. Journal of Cell Biology. 1999;147(2):307–320.

18. Karki S, Tokito M, Holzbaur E. A Dynactin Subunit with a Highly Conserved Cysteine-rich Motif Interacts Directly with Arp1. Journal of Biological Chemistry. 2000;275(7):4834–4839.

19. Garces J, Clark I, Meyer D, Vallee R. Interaction of the p62 subunit of dynactin with Arp1 and the cortical actin cytoskeleton. Current Biology. 1999;9(24):1497–1502.

20. Schroer T. DYNACTIN. Annual Review of Cell and Developmental Biology. 2004;20(1):759–779.

21. Urnavicius L, Zhang K, Diamant A, Motz C, Schlager M, Yu M et al. The structure of the dynactin complex and its interaction with dynein. Science. 2015;347(6229):1441–1446.

22. Cianfrocco M, DeSantis M, Leschziner A, Reck-Peterson S. Mechanism and Regulation of Cytoplasmic Dynein. Annual Review of Cell and Developmental Biology. 2015;31(1):83–108.

23. Echeverri C, Paschal B, Vaughan K, Vallee R. Molecular characterization of the 50-kD subunit of dynactin reveals function for the complex in chromosome alignment and spindle organization during mitosis. Journal of Cell Biology. 1996;132(4):617–633.

24. Salina D, Bodoor K, Eckley D, Schroer T, Rattner J, Burke B. Cytoplasmic Dynein as a Facilitator of Nuclear Envelope Breakdown. Cell. 2002;108(1):97–107.

25. Karki S, Holzbaur E. Cytoplasmic dynein and dynactin in cell division and intracellular transport. Current Opinion in Cell Biology. 1999;11(1):45–53.

26. Quintyne N, Gill S, Eckley D, Crego C, Compton D, Schroer T. Dynactin Is Required for Microtubule Anchoring at Centrosomes. Journal of Cell Biology. 1999;147(2):321–334.

27. Iyevleva A, Raskin G, Tiurin V, Sokolenko A, Mitiushkina N, Aleksakhina S et al. Novel ALK fusion partners in lung cancer. Cancer Letters. 2015;362(1):116–121.

28. Bransfield K, Askham J, Leek J, Robinson P, Mighell A. Phenotypic changes associated with DYNACTIN-2 (*DCTN2*) over expression characterise SJSA-1 osteosarcoma cells. Molecular Carcinogenesis. 2006;45(3):157–163.

29. Wang Q, Wang X, Liang Q, Wang S, Liao X, Li D et al. Prognostic Value of Dynactin mRNA Expression in Cutaneous Melanoma. Medical Science Monitor. 2018;24:3752–3763.

30. Wang S, Wang Q, Zhang X, Liao X, Wang G, Yu L et al. Distinct prognostic value of dynactin subunit 4 (*DCTN4*) and diagnostic value of *DCTN1, DCTN2*, and *DCTN4* in colon adenocarcinoma. Cancer Management and Research. 2018;Volume 10:5807–5824.

31. Schulze K, Imbeaud S, Letouzé E, Alexandrov L, Calderaro J, Rebouissou S et al. Exome sequencing of hepatocellular carcinomas identifies new mutational signatures and potential therapeutic targets. Nature Genetics. 2015;47(5):505–511.

32. Villanueva A, Newell P, Chiang D, Friedman S, Llovet J. Genomics and Signaling Pathways in Hepatocellular Carcinoma. Seminars in Liver Disease. 2007;27(1):055–076.

33. Tang G, Cho M, Wang X. OncoDB: an interactive online database for analysis of gene expression and viral infection in cancer. Nucleic Acids Research. 2022 Jan 7;50(D1):D1334–9.

34. Papatheodorou I, Fonseca NA, Keays M, Tang YA, Barrera E, Bazant W, Burke M, Füllgrabe A, Fuentes AM, George N, Huerta L. Expression Atlas: gene and protein expression across multiple studies and organisms. Nucleic acids research. 2018 Jan 4;46(D1):D246–51.

35. Wickham H, Chang W, Wickham MH. Package ‘ggplot2’. Create elegant data visualisations using the grammar of graphics. Version. 2016;2(1):1–89.

36. Allaire J. RStudio: integrated development environment for R. Boston, MA. 2012;770(394):165–71.

37. Pontén F, Jirström K, Uhlen M. The Human Protein Atlas—a tool for pathology. The Journal of Pathology: A Journal of the Pathological Society of Great Britain and Ireland. 2008 Dec;216(4):387–93.

38. Chandrashekar DS, Bashel B, Balasubramanya SA, Creighton CJ, Ponce-Rodriguez I, Chakravarthi BV, Varambally S. UALCAN: a portal for facilitating tumor subgroup gene expression and survival analyses. Neoplasia. 2017 Aug 1;19(8):649–58.

39. Goldman M, Craft B, Hastie M, Repečka K, McDade F, Kamath A, Banerjee A, Luo Y, Rogers D, Brooks AN, Zhu J. The UCSC Xena platform for public and private cancer genomics data visualization and interpretation. biorxiv. 2019 Jan 1:326470.

40. Liu CJ, Hu FF, Xia MX, Han L, Zhang Q, Guo AY. GSCALite: a web server for gene set cancer analysis. Bioinformatics. 2018 Nov 1;34(21):3771–2.

41. Gao J, Aksoy BA, Dogrusoz U, Dresdner G, Gross B, Sumer SO, Sun Y, Jacobsen A, Sinha R, Larsson E, Cerami E. Integrative analysis of complex cancer genomics and clinical profiles using the cBioPortal. Science signaling. 2013 Apr 2;6(269):pl1-.

42. Gentles AJ, Newman AM, Liu CL, Bratman SV, Feng W, Kim D, Nair VS, Xu Y, Khuong A, Hoang CD, Diehn M. The prognostic landscape of genes and infiltrating immune cells across human cancers. Nature medicine. 2015 Aug;21(8):938–45.

43. Tang Z, Kang B, Li C, Chen T, Zhang Z. GEPIA2: an enhanced web server for large-scale expression profiling and interactive analysis. Nucleic acids research. 2019 Jul 2;47(W1):W556–60.

44. Li T, Fu J, Zeng Z, Cohen D, Li J, Chen Q, Li B, Liu XS. TIMER2. 0 for analysis of tumor-infiltrating immune cells. Nucleic acids research. 2020 Jul 2;48(W1):W509–14.

45. Racle J, Gfeller D. EPIC: a tool to estimate the proportions of different cell types from bulk gene expression data. InBioinformatics for Cancer Immunotherapy 2020 (pp. 233–248). Humana, New York, NY.

46. Yu G, Wang LG, Han Y, He QY. clusterProfiler: an R package for comparing biological themes among gene clusters. Omics: a journal of integrative biology. 2012 May 1;16(5):284–7.

47. Tsuchiya M, Parker JS, Kono H, Matsuda M, Fujii H, Rusyn I. Gene expression in nontumoral liver tissue and recurrence-free survival in hepatitis C virus-positive hepatocellular carcinoma. Molecular cancer. 2010 Dec;9(1):1–1.

48. Diaz G, Engle RE, Tice A, Melis M, Montenegro S, Rodriguez-Canales J, Hanson J, Emmert-Buck MR, Bock KW, Moore IN, Zamboni F. Molecular Signature and Mechanisms of Hepatitis D Virus–Associated Hepatocellular CarcinomaGenomics of HDV-HCC Suggests Potential Drivers. Molecular Cancer Research. 2018 Sep 1;16(9):1406–19.

49. Gentleman RC, Carey VJ, Bates DM, Bolstad B, Dettling M, Dudoit S, Ellis B, Gautier L, Ge Y, Gentry J, Hornik K. Bioconductor: open software development for computational biology and bioinformatics. Genome biology. 2004 Sep;5(10):1–6.

50. Smyth GK. Limma: linear models for microarray data. InBioinformatics and computational biology solutions using R and Bioconductor 2005 (pp. 397–420). Springer, New York, NY.

51. Liang P, Pardee AB. Analysing differential gene expression in cancer. Nature Reviews Cancer. 2003 Nov;3(11):869–76.

52. Liñares Blanco J, Gestal M, Dorado J, Fernandez-Lozano C. Differential gene expression analysis of RNA-seq data using machine learning for Cancer research. InMachine Learning Paradigms 2019 (pp. 27–65). Springer, Cham.

53. Duraiyan J, Govindarajan R, Kaliyappan K, Palanisamy M. Applications of immunohistochemistry. Journal of pharmacy & bioallied sciences. 2012 Aug;4(Suppl 2):S307.

54. Kulis M, Esteller M. DNA methylation and cancer. Advances in genetics. 2010 Jan 1;70:27–56.

55. Luczak MW, Jagodziński PP. The role of DNA methylation in cancer development. Folia histochemica et cytobiologica. 2006;44(3):143–54.

56. Koch A, Joosten SC, Feng Z, de Ruijter TC, Draht MX, Melotte V, Smits KM, Veeck J, Herman JG, Van Neste L, Van Criekinge W. Analysis of DNA methylation in cancer: location revisited. Nature reviews Clinical oncology. 2018 Jul;15(7):459–66.

57. Zhu JD. The altered DNA methylation pattern and its implications in liver cancer. Cell research. 2005 Apr;15(4):272–80.

58. Bai Y, Tong W, Xie F, Zhu L, Wu H, Shi R, Wang L, Yang L, Liu Z, Miao F, Zhao Q. DNA methylation biomarkers for diagnosis of primary liver cancer and distinguishing hepatocellular carcinoma from intrahepatic cholangiocarcinoma. Aging (Albany NY). 2021 Jul 15;13(13):17592.

59. Zhang YJ, Wu HC, Shen J, Ahsan H, Tsai WY, Yang HI, Wang LY, Chen SY, Chen CJ, Santella RM. Predicting hepatocellular carcinoma by detection of aberrant promoter methylation in serum DNA. Clinical Cancer Research. 2007 Apr 15;13(8):2378–84.

60. Sceusi EL, Loose DS, Wray CJ. Clinical implications of DNA methylation in hepatocellular carcinoma. Hpb. 2011 Jun 1;13(6):369–76.

61. Worm J, Guldberg P. DNA methylation: an epigenetic pathway to cancer and a promising target for anticancer therapy. Journal of oral pathology & medicine. 2002 Sep;31(8):443–9.

62. Martincorena I, Campbell PJ. Somatic mutation in cancer and normal cells. Science. 2015 Sep 25;349(6255):1483–9.

63. Iranzo J, Martincorena I, Koonin EV. Cancer-mutation network and the number and specificity of driver mutations. Proceedings of the National Academy of Sciences. 2018 Jun 26;115(26):E6010–9.

64. Shao X, Lv N, Liao J, Long J, Xue R, Ai N, Xu D, Fan X. Copy number variation is highly correlated with differential gene expression: a pan-cancer study. BMC medical genetics. 2019 Dec;20(1):1–4.

65. Guo L, Jing Y. Construction and identification of a novel 5-gene signature for predicting the prognosis in breast cancer. Frontiers in Medicine. 2021;8.

66. Rao CV, Asch AS, Yamada HY. Frequently mutated genes/pathways and genomic instability as prevention targets in liver cancer. Carcinogenesis. 2017 Jan 1;38(1):2–11.

67. Garnelo M, Tan A, Her Z, Yeong J, Lim CJ, Chen J, Lim KH, Weber A, Chow P, Chung A, Ooi LL. Interaction between tumour-infiltrating B cells and T cells controls the progression of hepatocellular carcinoma. Gut. 2017 Feb 1;66(2):342–51.

68. Floudas CS, Brar G, Greten TF. Immunotherapy: current status and future perspectives. Digestive Diseases and Sciences. 2019 Apr;64(4):1030–40.

69. Anuraga G, Tang WC, Phan NN, Ta HD, Liu YH, Wu YF, Lee KH, Wang CY. Comprehensive Analysis of Prognostic and Genetic Signatures for General Transcription Factor III (GTF3) in clinical colorectal cancer patients using bioinformatics approaches. Current Issues in Molecular Biology. 2021 Apr 27;43(1):2–0.

70. Mo X, Li T, Xie Y, Zhu L, Xiao B, Liao Q, Guo J. Identification and functional annotation of metabolism□associated lnc RNA s and their related protein□coding genes in gastric cancer. Molecular Genetics & Genomic Medicine. 2018 Sep;6(5):728–38.

71. Elizalde M, Urtasun R, Azkona M, Latasa MU, Goñi S, García-Irigoyen O, Uriarte I, Segura V, Collantes M, Di Scala M, Lujambio A. Splicing regulator SLU7 is essential for maintaining liver homeostasis. The Journal of clinical investigation. 2014 Jul 1;124(7):2909–20.

72. Zeng Y, Tan X, Gong J, Liu Z. Exosome-mediated miR-1290 regulation of SLU7 affects hepatocellular carcinoma process. SSRN Preprint. 2020

73. Lv M, Cao D, Zhang L, Hu C, Li S, Zhang P, Zhu L, Yi X, Li C, Yang A, Yang Z. METTL9 mediated N1-histidine methylation of zinc transporters is required for tumor growth. Protein & cell. 2021 Dec;12(12):965–70.

74. Cheng Y, Liu W, Kim ST, Sun J, Lu L, Sun J, Zheng SL, Isaacs WB, Xu J. Evaluation of PPP2R2A as a prostate cancer susceptibility gene: a comprehensive germline and somatic study. Cancer genetics. 2011 Jul 1;204(7):375–81.

75. Liang WL, Cao J, Xu B, Yang P, Shen F, Sun Z, Li WL, Wang Q, Liu F. miR-892a regulated PPP2R2A expression and promoted cell proliferation of human colorectal cancer cells. Biomedicine & Pharmacotherapy. 2015 May 1;72:119–24.

76. Beca F, Pereira M, Cameselle-Teijeiro JF, Martins D, Schmitt F. Altered PPP2R2A and Cyclin D1 expression defines a subgroup of aggressive luminal-like breast cancer. BMC cancer. 2015 Dec;15(1):1–0.

77. Sarkar S, Horn G, Moulton K, Oza A, Byler S, Kokolus S, Longacre M. Cancer development, progression, and therapy: an epigenetic overview. International journal of molecular sciences. 2013 Oct 21;14(10):21087–113.

78. Aird KM, Zhang R. Nucleotide metabolism, oncogene-induced senescence and cancer. Cancer letters. 2015 Jan 28;356(2):204–10.

79. Albert M, Helin K. Histone methyltransferases in cancer. InSeminars in cell & developmental biology 2010 Apr 1 (Vol. 21, No. 2, pp. 209–220). Academic Press.

80. Kim JA. Peroxisome metabolism in cancer. Cells. 2020 Jul 14;9(7):169.

