## Supplementary Information for "A Multi-omics Study on the Oncogenic Roles and Clinical Significance of Dynactin Family Gene (*DCTN1-6*) Expression in Liver Hepatocellular Carcinoma"

| **Sl. No.** | **Features** | **Comparison** | **p-value** | | | | | |
| --- | --- | --- | --- | --- | --- | --- | --- | --- |
|  |  |  | **DCTN1** | **DCTN2** | **DCTN3** | **DCTN4** | **DCTN5** | **DCTN6** |
| **1.** | **Age** | Normal-vs-Age (21-40 Yrs) | 3.77E-05 | 3.54E-08 | 1.40E-03 | 3.16E-06 | 2.74E-06 | **9.89E-02** |
|  |  | Normal-vs-Age (41-60 Yrs) | <1E-12 | <1E-12 | <1E-12 | 1.62E-12 | 1.64E-12 | 9.63E-04 |
|  |  | Normal-vs-Age (61-80 Yrs) | <1E-12 | 1.62E-12 | 1.62E-12 | <1E-12 | 1.69E-12 | **2.86E-01** |
|  |  | Normal-vs-Age (81-100 Yrs) | 1.06E-02 | 1.84E-03 | 1.51E-02 | 1.19E-02 | 7.85E-02 | **7.14E-01** |
| **2.** | **Individual Cancer Stage** | Normal-vs-Stage1 | 1.11E-16 | <1E-12 | 1.62E-12 | 1.62E-12 | 1.65E-12 | **1.21E-01** |
|  |  | Normal-vs-Stage2 | 3.64E-14 | 1.06E-14 | 9.21E-14 | 7.32E-15 | 2.01E-10 | **1.40E-01** |
|  |  | Normal-vs-Stage3 | 1.62E-12 | <1E-12 | 1.62E-12 | 1.62E-12 | 5.05E-11 | 7.35E-04 |
|  |  | Normal-vs-Stage4 | 2.41E-02 | 2.75E-02 | 3.91E-02 | **8.87E-02** | **2.11E-01** | 1.34E-02 |
| **3.** | **Tumor Grade** | Normal-vs-Grade1 | 6.66E-08 | 6.00E-12 | 3.42E-10 | 1.45E-10 | 1.04E-03 | **5.26E-01** |
|  |  | Normal-vs-Grade2 | 1.62E-12 | <1E-12 | 1.62E-12 | 1.62E-12 | 1.64E-12 | 4.12E-02 |
|  |  | Normal-vs-Grade3 | <1E-12 | <1E-12 | 1.62E-12 | <1E-12 | 1.62E-12 | 7.18E-04 |
|  |  | Normal-vs-Grade4 | 1.67E-03 | 3.77E-04 | 2.28E-02 | 6.56E-03 | 5.20E-03 | **7.42E-01** |
| **4.** | **Nodal Metastasis Status** | Normal-vs-N0 | <1E-12 | <1E-12 | 1.62E-12 | <1E-12 | 1.62E-12 | 8.83E-04 |
|  |  | Normal-vs-N1 | **5.00E-02** | **9.20E-02** | 2.75E-02 | 1.62E-12 | **8.58E-02** | **1.75E-01** |

**Supplementary Table S1:** The result of association analysis between *DCTN1-6* expression and LIHC patients’ clinicopathological characteristics. All the *DCTNs* except *DCTN6* showed noticeable association with most of the clinical parameters examined. The hypothesis was evaluated by performing a student’s *t-test* across TCGA LIHC cohorts through the UALCAN server.

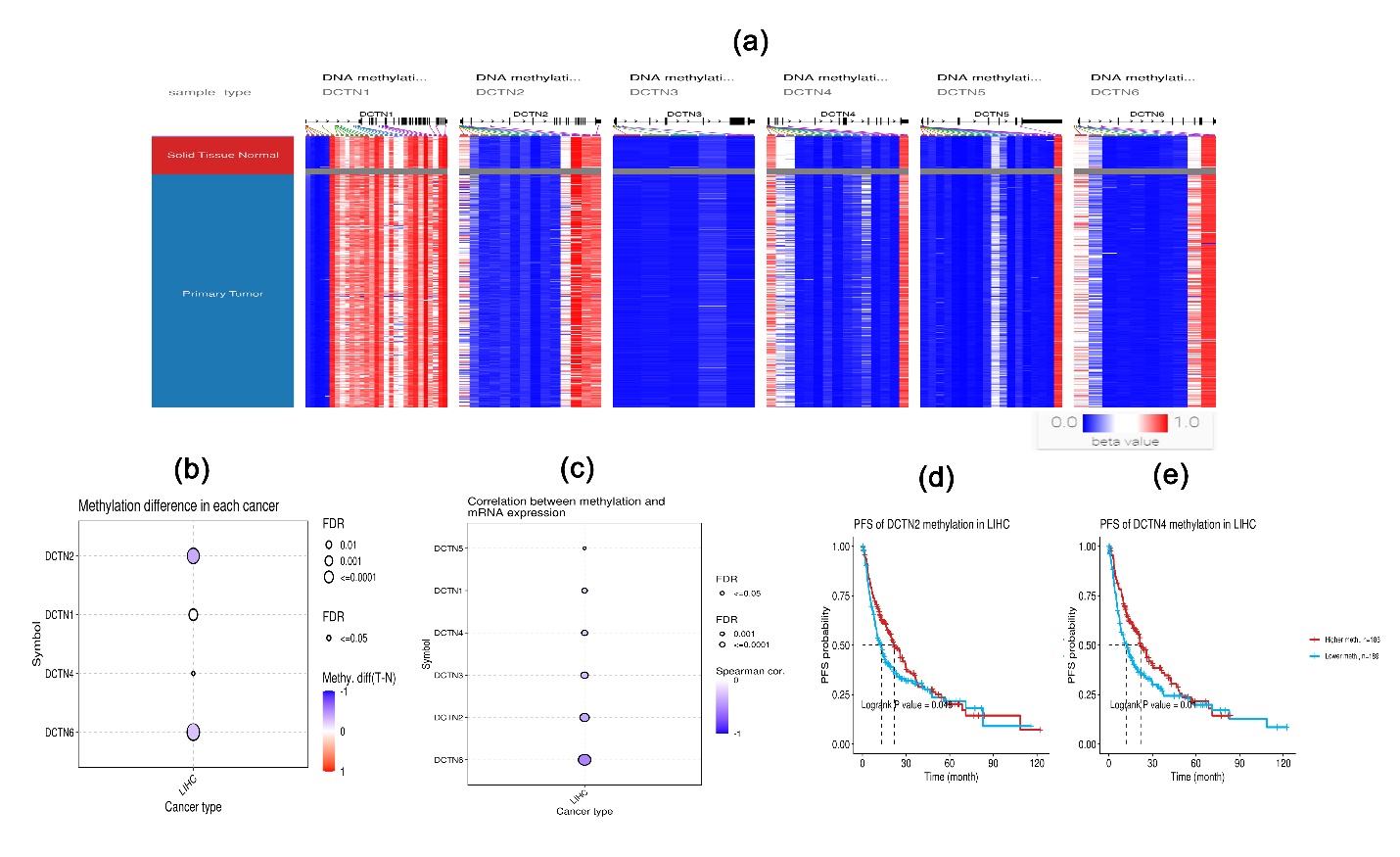

**Supplementary Figure S1:** The DNA methylation pattern of *DCTN* coding genes in TCGA LIHC samples established by inspecting the methylation 450k Illumina sequencing data (a). The grey region refers to the lack of methylation data in accordance with the sample type represented in the leftmost column. The methylation differences in *DCTN* coding genes between normal liver and LIHC tissues (b). Correlation between *DCTN* methylation and their mRNA expression level in LIHC tissues (c). The association of LIHC patients’ PFS with *DCTN2* (d) and *DCTN4* (e) methylation. Illumina beta value identifier: 0: unmethylated; 0.5: hemimethylated (50% methylation); 1: fully methylated.

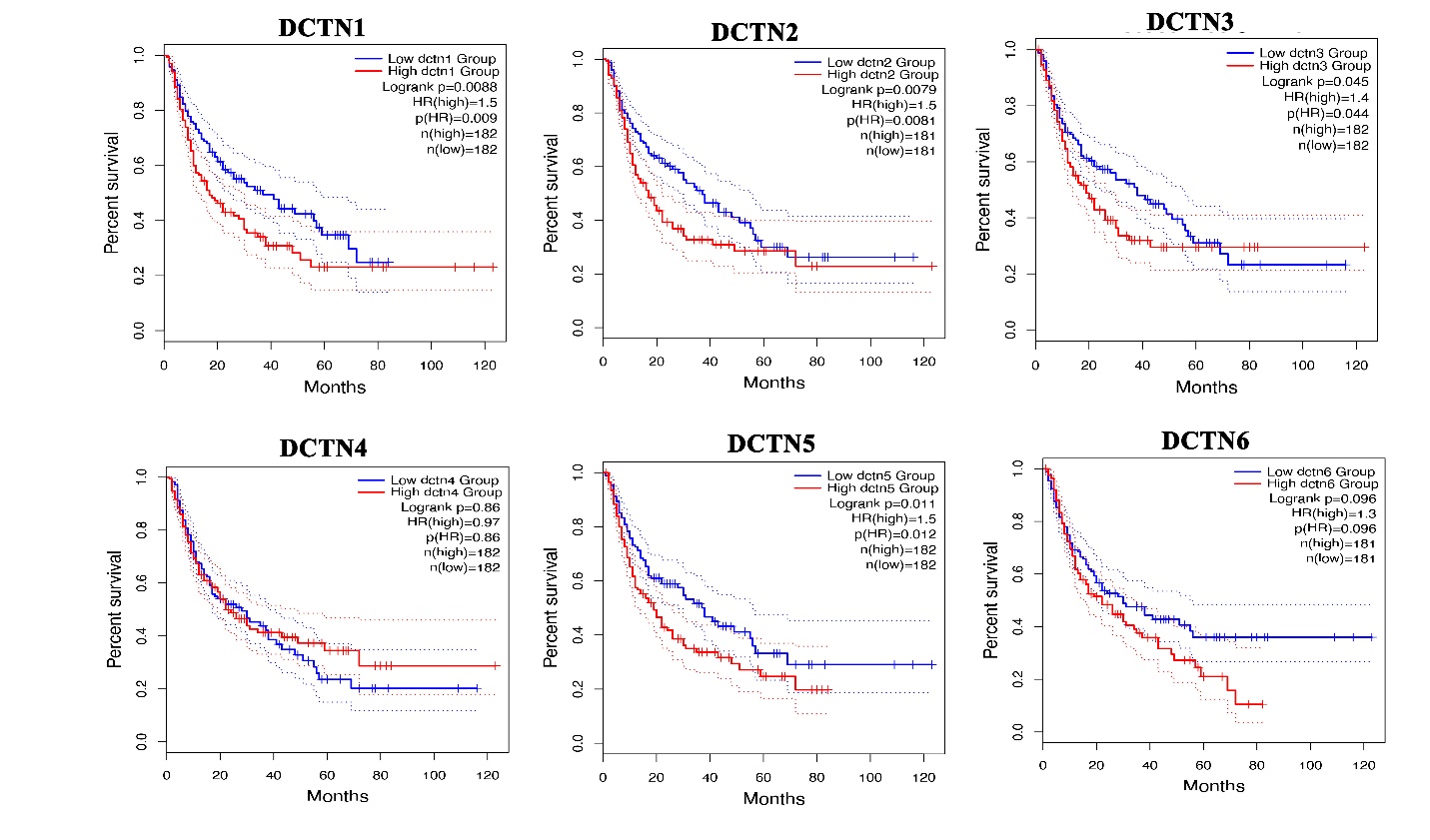

**Supplementary Figure S2:** KM plot representation of the association between *DCTN* family gene expression and LIHC patients’ RFS. The red and blue plots represent the higher and lower *DCTN* expressed by LIHC patients, respectively. The vertical mark along the plot indicates an event (death). Overexpression of the *DCTNs* was found to be significantly associated (except for *DCTN4* and *DCTN6*) with the poor RFS of LIHC patients.

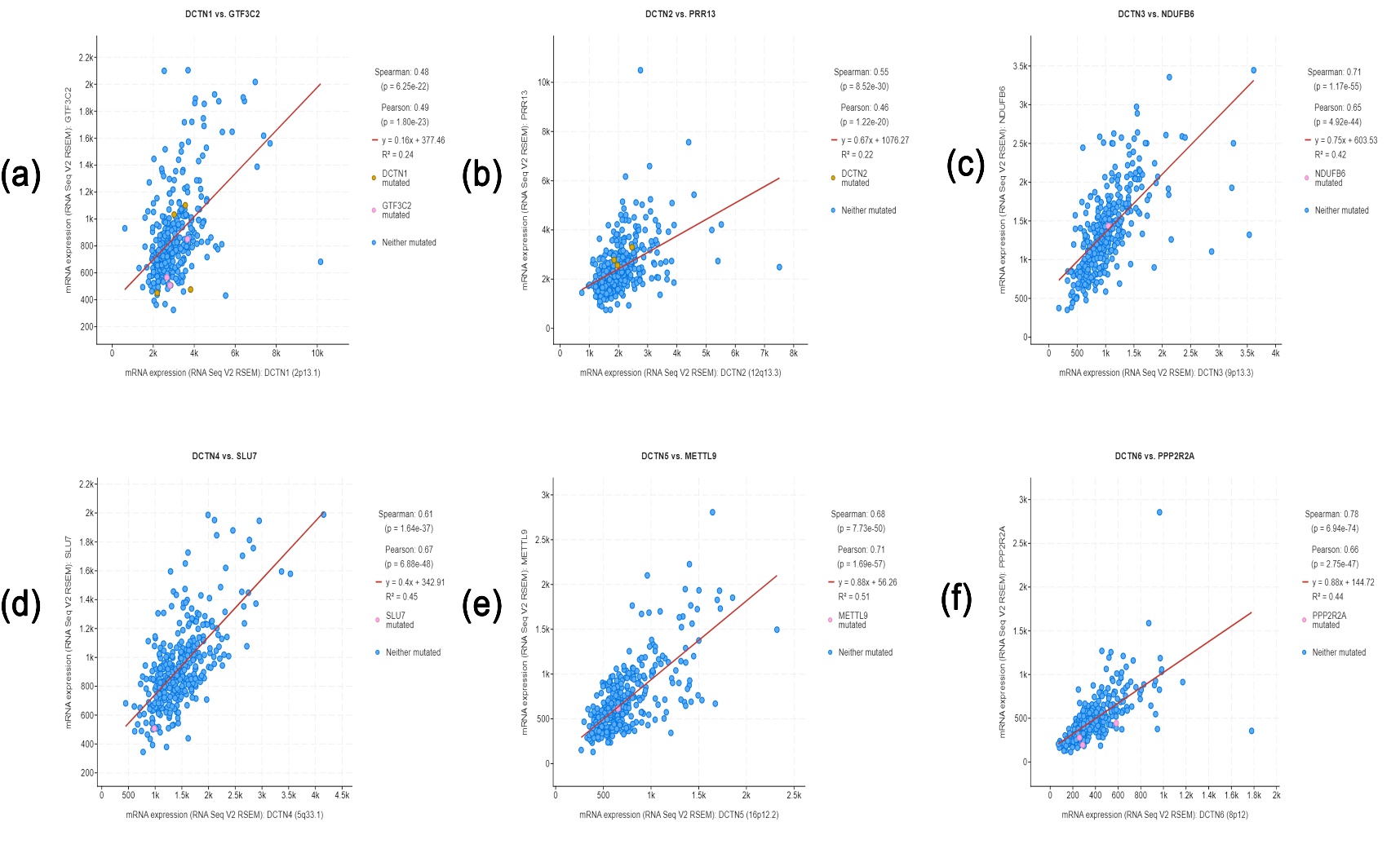

**Supplementary Figure S3:** Scatterplot representation of the top positively co-expressed genes of (a) *DCTN1*, (b) *DCTN2*, (c) *DCTN3*, (d) *DCTN4*, (e) *DCTN5*, (f) *DCTN6* in LIHC tissues obtained from the TCGA LIHC study (Firehose Legacy) in cBioPortal server. RNA-seq V2 RSEM normalized scores of the sequencing reads were compared to the expression values. RSEM: RNA-seq by Expectation-Maximization.

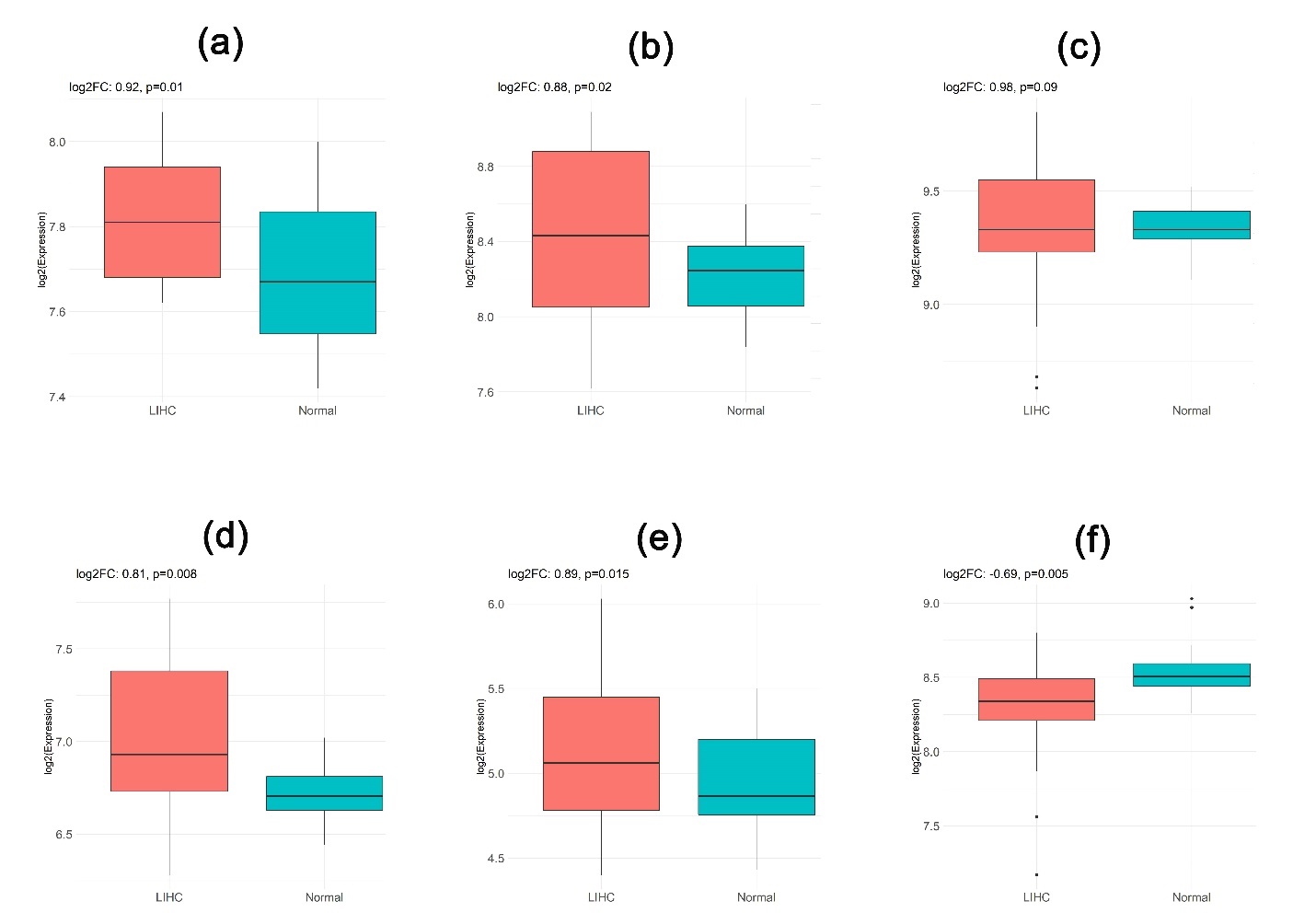

**Supplementary Figure S4:** The expression pattern of *DCTNs* in LIHC tissues (n=16) and adjacent normal liver tissues (n=28) from GSE98383 microarray dataset: (a) *DCTN1*, (b) *DCTN2*, (c) *DCTN3*, (d) *DCTN4*, (e) *DCTN5*, (f) *DCTN6*. All the genes except *DCTN6* were found to be overexpressed in LIHC tissues.
